# Sensitive tumor detection, accurate quantification, and cancer subtype classification using low-pass whole methylome sequencing of plasma DNA

**DOI:** 10.1101/2024.06.10.598204

**Authors:** Marta Paoli, Francesca Galardi, Agostina Nardone, Chiara Biagioni, Dario Romagnoli, Samantha Di Donato, Gian Marco Franceschini, Luca Livraghi, Marta Pestrin, Giuseppina Sanna, Emanuela Risi, Ilenia Migliaccio, Erica Moretti, Luca Malorni, Laura Biganzoli, Francesca Demichelis, Matteo Benelli

## Abstract

The analysis of circulating tumor DNA (ctDNA) is increasingly used for monitoring disease in patients with metastatic cancer. Here, we introduce a robust and reproducible strategy combining low-pass whole methylome sequencing of plasma DNA with METER, a novel computational tool. Engaging prediction models trained on independent available datasets, METER enables the detection and quantification of tumor content (TC) and performs molecular cancer subtyping. Applied to plasma methylomes from early metastatic breast cancer patients, our method demonstrated reliable quantification, sensitive tumor detection below 3% of TC, and the ability to perform accurate Estrogen Receptor (ER) subtyping. METER provided clinically relevant predictions, underscored by associations with relevant prognostic factors, robust correlation with matched circulating tumor cells, and highly correlated with patients’ outcomes in challenging scenarios as TC<3%. Our strategy provides comprehensive and sensitive analysis of plasma samples, serving as a valuable yet cost-effective precision oncology tool.

## Introduction

The assessment of cell-free tumor DNA in the bloodstream of cancer patients (circulating tumor DNA, ctDNA) is becoming critical for the clinical management of oncological patients (*1, 2*). Based on a large body of clinical studies, it is expected that liquid biopsy will play a significant role in clinical practice in the near future. In the metastatic setting, liquid biopsy can complement imaging to improve the monitoring of treatment response (*3*), and presents several advantages over traditional tissue biopsies often limited by challenging-to-access metastatic sites. The minimal invasiveness of liquid biopsy reduces its impact on patients and allows for serial sampling, overall increasing the capability for real-time disease monitoring and timely interventions in treatment strategies (*4*).

The presence and the amount of ctDNA (Tumor Content, TC) has been shown to be prognostically relevant and associated with early progression in many cancers, including breast cancer (BC) (*5, 6*). One of the most common and effective strategy for measuring TC is through the analysis of Copy Number Alterations (CNA) either from target sequencing (*7, 8*) or low-pass Whole Genome Sequencing (lpWGS) by ichorCNA (*9*) . This approach has several advantages compared to other common proposed methods based on the analysis of circulating mutations (*10*), methylomics (*11*), and fragmentomics (*12*) . Indeed, it requires low input DNA amount, enhancing the likelihood of successful analysis of liquid biopsy samples. Second, it is tumor type agnostic, requiring no prior information about the molecular characteristics of the metastases shedding DNA in the circulation. Third, it is highly reproducible and cost effective as it relies on well-established library preparation protocols, requires less than 30 million reads, and utilizes state-of-the-art computational tools. However, the typical lower limit of TC detection of ∼3% makes it unsuitable to applications as longitudinal monitoring where the detection of TC during the initial cycles of treatment is critical, especially for early metastatic lines (*13*). Further, low-pass CNA based approaches fail to offer information about tumor subtypes, typically characterized by specific transcriptional programs that cannot be easily estimated by genomic alterations. Recent methods based on the analysis of DNA-methylation patterns (*11*), epigenomics profiling (*14, 15*) and nucleosome positioning (*16*) enable the analysis of relevant molecular features typically measurable in tumor tissues, such as histological and molecular subtypes. However, most of these epigenetics-based approaches require non-trivial experimental procedures, deep sequencing, prior knowledge on epigenetics features and assistance of machine learning classifiers for their application.

In this study, we introduce a cost-effective and accurate strategy that offers, in a single approach, both detection and quantification of TC and accurate estimation of phenotypic subtyping in circulation. Our approach is based on the analysis of low-pass Whole Genome Bisulfite Sequencing (lpWGBS), using a novel computational tool called METER (DNA-<u>MEt</u>hylome Analys<u>ER</u>). METER incorporates strategies for the comprehensive analysis of tumor signal in circulation, including quantification, detection and subtyping. To evaluate METER’s performance, we generated a lpWGBS dataset of the the MIMESIS cohort (*17*), including 124 plasma samples from 59 patients with metastatic BC (mBC). Specifically, the MIMESIS dataset represents a challenging scenario for the analysis of TC in the metastatic setting, as 68% of patients had received less than two lines of metastatic treatment and are therefore expected to represent low TC states. Leveraging curated clinical data from the MIMESIS study, we evaluated the utility of METER in providing clinically relevant information by examining its association with other liquid biopsy analysis, clinical characteristics, and outcomes.

## Results

### The MIMESIS study

To fill the gap of plasma TC quantification and detection in the low TC range, relevant to non-invasive disease monitoring of early metastatic disease, we reasoned that identifying robust tumor-specific DNA-methylation sites and regions through comparative analysis of the BC BASIS (*18*) cohort and healthy cell-free DNA (*19*) WGBS data would be instrumental. The qualification cohort consists of plasma samples from patients with mBC collected within the Minimal DNA-methylation Signatures (MIMESIS) study specifically designed for the assessment of optimal experimental and computational strategies for the analysis of TC For this specific study, lpWGBS of cell-free DNA of 124 plasma samples from 59 patients from the MIMESIS program and 30 samples from healthy donors was generated. mBC blood drowns were collected before starting the treatment with systemic therapy (pre-treatment baseline, BL; N=59), after the 1^st^ cycle of treatment (day 1 of cycle 2, C2D1; N=43), and at progression (Prog; N=22), respectively. A total of 45 out of 59 (76%) and 14 out of 59 (24%) patients were ER+ and ER-, respectively. At study entry, 88% of patients (52 out of 59) had undergone less than 3 metastatic treatment lines (range 0-6 previous lines). Furthermore, 90% of patients (53 out of 59) had fewer than 4 metastatic sites at study entry, while the remaining patients presented a maximum of 6 metastatic sites. A per-sample median coverage of 0.8 (range 0.3-2.0) was obtained, corresponding to a median of 21 (range 7-50) million reads per sample. Main clinical characteristics of patients included in the MIMESIS lpWGBS cohort are reported in **Table S1**.

### The METER computational tool

METER is a computational tool composed of three main modules (**Fig. 1B**): i) METER-quant, to measure TC, ii) METER-detect, to detect the presence (METER+) or absence (METER-) of ctDNA, and iii) METER-subtype, to infer molecular subtype. Briefly, each module relies on task-specific informative Differentially Methylated Regions (iDMR) and Sites (iDMS) identified through the differential analysis between DNA-methylation data from tumor-type specific tissue samples and control samples such as cell-free DNA (cfDNA) samples from healthy donors or whole blood. DMR and DMS are filtered based on different, module-specific stringent criteria including beta difference, AUC and beta distribution of control samples. The final list including passing filter DMR (iDMR) and DMS (iDMS) are used differently in the three modules. In this study focusing on liquid biopsies from patients with mBC, Rocker-meth was used to detect DMR and DMS comparing WGBS of BC tumors from the BASIS dataset (*18*) and healthy cell-free DNA from the Fox-Fisher et al. dataset(*19*). The METER-subtype module was applied to infer Estrogen Receptor (ER) status from the circulation. Detailed illustration of each module is depicted in **Fig. 1C** and descripted in **Material and Methods**.

**Fig. 1.**
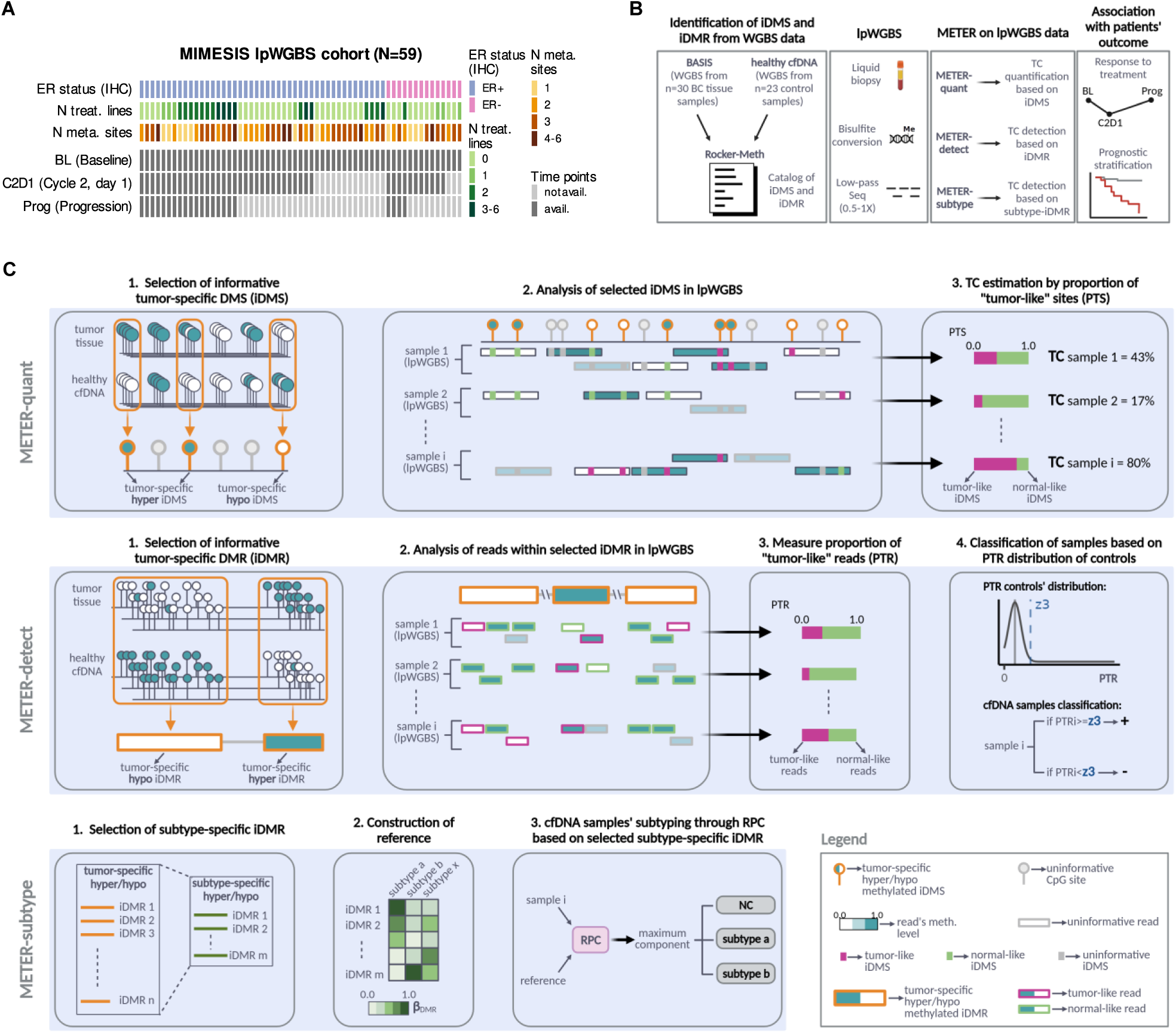
Study cohort and METER tool. **(A)** Heatmap showing the MIMESIS lpWGBS study cohort comprising 124 samples collected at multiple time points (BL, C2D1, Prog) from 59 patients with mBC. For each patient availability of samples at different time points along with relevant clinical characteristics is reported. **(B)** Schematic of the study showing (from left to right): identification of iDMS and iDMR, generation of lpWGBS of liquid biopsy samples, application of METER computational tool and its modules, association of METER results with patients’ clinical data. **(C)** Overview of the METER computational tool. METER is a computational tool to analyse TC, exploiting tumor-specific iDMS and iDMR in lpWGBS data, comprising 3 modules: METER-quant (top panel) to measure ctDNA in circulation; METER-detect (middle panel) to classify samples as ctDNA +/-; METER-subtype (bottom panel) to infer tumor subtype in circulation. Each module relies on task-specific iDMS and iDMR. lpWGBS, low-pass whole genome bisulfite sequencing; IHC, immunohistochemistry; N treat. lines, number of metastatic treatment lines at study entry; N meta. sites, number of metastatic sites at study entry; BL, baseline; C2D1, cycle 2 day 1; Prog, progression of disease; iDMS, informative differentially methylated sites; iDMR, informative differentially methylated regions; mBC, metastatic breast cancer; ctDNA, circulating tumor DNA; β_DMR_:beta by DMR; RPC: robust partial correlation; NC: not classified.

### Tumor content quantification by METER-quant

Obtaining an accurate estimation of TC, the fraction of ctDNA over the total cfDNA shed into the bloodstream is emerging as a crucial clinical need. Numerous studies have highlighted its role as an independent prognostic factor in various cancers, including breast cancer (BC) (*5*). Elevated levels of TC are typically associated with relevant molecular features such as actionable variants and high mutational and copy number burden (*20, 21*). Accurate quantification of TC is also essential for inferring the clonal status of somatic mutations, particularly in monitoring the efficacy of targeted therapies and understanding emerging molecular mechanisms of resistance (*7*). We hypothesized the existence of prevalent and pervasive, tumor-type-specific iDMS (**Table S2)** capable of providing precise quantification of TC and implemented this strategy in METER-quant exploiting the proportion of tumor-like sites (PTS) (**Online Methods**). To evaluate the performance of METER-quant, we therefore applied it to 124 plasma samples of the MIMESIS lpWGBS dataset (**Table S3**). In tumor samples, TC ranged between 0 and 52% and was statistically significantly higher (p<1e-10) than TC values in controls, as expected (**Fig. 2A**). We then compared METER-quant estimates with those from ichorCNA (*9*) applied to the same data. Since ichorCNA was originally designed for lpWGS, not lpWGBS, we performed a preliminary study to ensure the applicability of ichorCNA in this type of data. Using a cohort of 9 cfDNA samples from patients with advanced metastatic prostate cancer (*22*), we found excellent concordance between CNA profiles derived from lpWGBS using ichorCNA and those from matched high-coverage WES (200X). Furthermore, TC estimates obtained by ichorCNA were in line with reference estimates obtained by an independent CNA based method (*23*), overall indicating that bisulfite treatment does not introduce significant bias in CNA estimation and therefore does not hinder ichorCNA applicability in these data (**Supplementary Materials and Fig. S1** and **S2**). The TC values for the 124 cfDNA mBC samples estimated by METER-quant were highly concordant with those obtained from the state-of-the-art tool ichorCNA (R>0.90, p<1e-50) (**Fig. 2B**). In particular, METER-quant showed accurate and comparable estimation also for low TC values, in the range of 3%-10% as estimated by ichorCNA. For lower values, we observed poor concordance between the two methods (R=0.27, p=0.04), as expected based on the inaccuracy of ichorCNA’s TC estimations due to its lower limit of detection of 3%. To examine the ability of METER-quant to provide accurate measures for low-level TC, we generated artificial samples consisting of serial dilutions (0%, 0.1%, 0.5%, 1%, 2.5%, 5%) of DNA from luminal T47D BC cell line culture medium, as a surrogate of ctDNA, with commercial human genomic DNA from white blood cells (**Fig. 2C**). As expected, we observed that ichorCNA loose linearity in predicting TC (Median Absolute Deviation, MAE=0.01, Root Mean Square Deviation RMSE=0.01), still maintaining moderately high correlation (R=0.95, rho=0.86) with expected values. On the other hand, METER-quant showed accurate estimation down to 0.5-1%, obtaining higher correlation with expected values (R=0.96, rho=0.95) and lower dispersion (MAE=0.005, RMSE=0.006) than ichorCNA (**Fig. 2C** and **Table S4**). We finally tested whether the use of a model for training METER-quant could enhance the accuracy of TC estimations. We therefore developed a linear model using 60% of lpWGBS samples from our study cohort as a training set (N=94, including 18 controls and 76 tumors), while the remaining 40% (N=60) served as a test set (model parameters reported in **Table S5)**. PTS was used as the predictor variable, and observed values were used as references (ichorCNA TC estimates for tumor samples and 0% TC for control samples). Within the test set, control samples exhibited a median TC of 0% (range 0-1%), whereas tumor samples had a significantly higher median TC of 4% (range 0-60%), compared to controls (p<1e-5) and a strong concordance between observed and predicted values within the test set (R=0.94, p<1e-28) was obtained (**Fig. S3**).

**Fig. 2.**
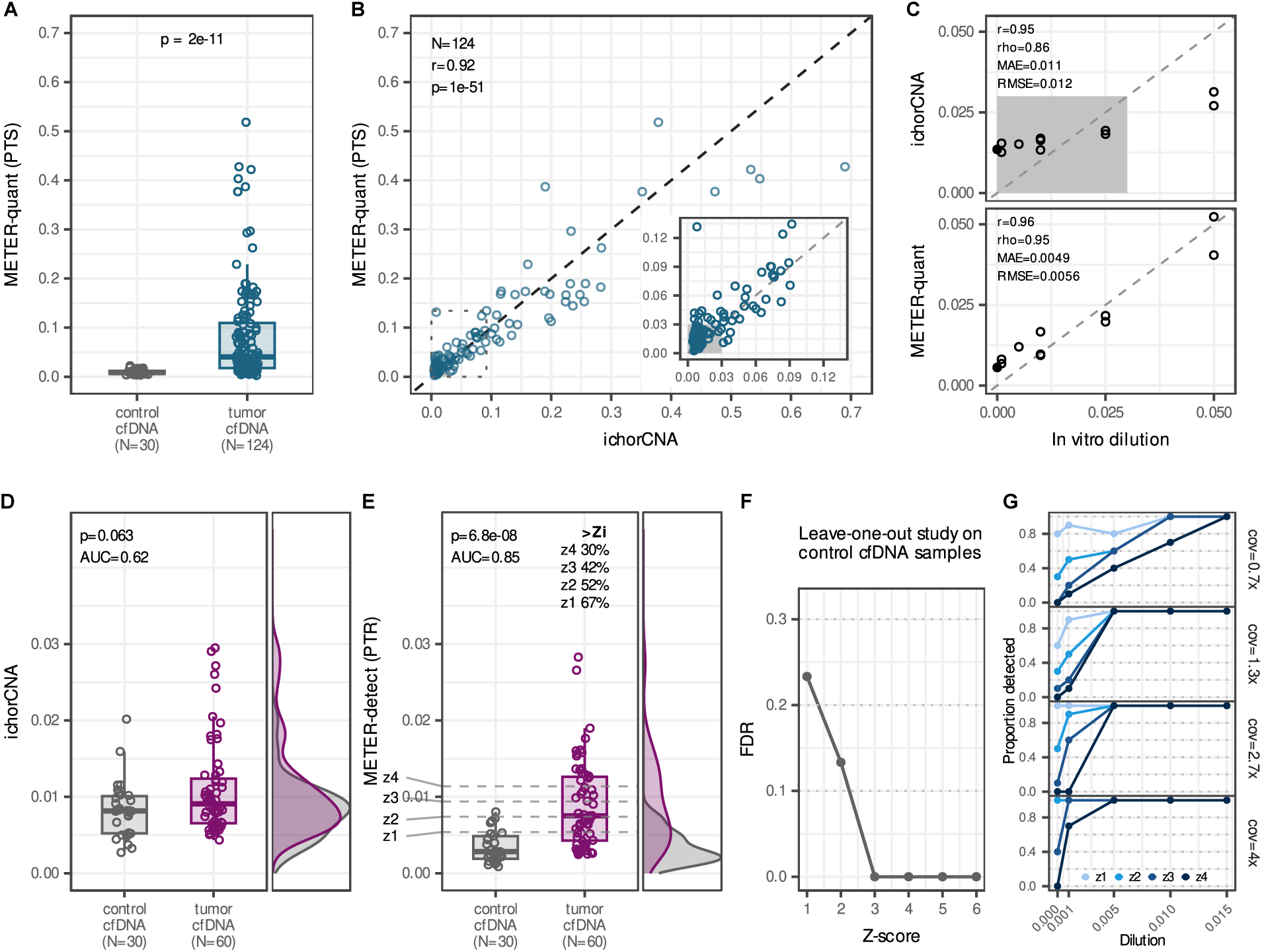
TC quantification by METER-quant and TC detection by METER-detect. **(A)** Boxplots showing the distribution of TC estimates by METER-quant in control and mBC cfDNA samples within the study cohort. P-value was estimated by Wilcoxon test **(B)** Scatter plot showing TC estimates of mBC cfDNA samples by METER-quant (y-axis) and ichorCNA (x-axis). The inset plot reports samples with TC range of 0-10% according to ichorCNA. The shaded region highlights the range of TC where ichorCNA is not applicable (0-3%). r and p are Pearson’s correlation coefficient and corresponding p-value, respectively **(C)** Scatter plots of TC by ichorCNA (top panel) and METER-quant (bottom panel) versus in vitro serial dilutions of DNA from T47D BC cell line culture medium with commercial human genomic DNA from white blood cells. Filled dot represents 0% TC dilution. The shaded region refers to the range of TC where ichorCNA is not applicable (0-3%). Boxplots and density plots showing the distribution of TC by ichorCNA (**D**) and PTR by METER-detect (**E**) in 30 control and 60 mBC cfDNA samples within the study cohort with TC by ichorCNA<3%. In E, the dashed lines indicate different Z-score thresholds based on control samples’ PTR distribution and percentages of tumor samples with Z-scores exceeding Z-score thresholds (Z_i_) are shown. p is p-value estimated by Wilcoxon test. **(F)** Leave-one-out study on control cfDNA samples to estimate FDR of METER-detect at different Z-score thresholds. **(G)** Line-plots showing in-silico dilutions study to evaluate detection sensitivity and FDR of METER-detect at increasing mean coverage levels (top to bottom panels) and Z-score thresholds. PTS, proportion of tumor-like sites; r: Pearson’s correlation coefficient; rho: Spearman’s correlation coefficient; MAE: mean absolute error; RMSE: root mean squared error; PTR: proportion of tumor-like reads; AUC: area under the curve; FDR: false discovery rate; TC: tumor content; mBC: metastatic breast cancer; cfDNA: cell-free DNA.

### Tumor detection by METER-detect

The accurate detection of the presence of tumor DNA in the bloodstream of oncological patients, regardless of TC quantification, provides classification of patients into ctDNA positive or negative, serving as a pragmatic strategy to obtain clinically relevant stratification of patients and as a tool for guiding treatment decisions. We therefore developed METER-detect, a computational strategy exploiting specific iDMR (**Table S6**) for the detection of tumor signal in circulation from lpWGBS and implemented this method into the METER tool. To test the ability of METER-detect in classifying samples based on the presence (ctDNA positive, or METER+) or the absence (ctDNA negative, or METER-) of tumor DNA in circulation, we focused on 60 samples that scored below ichorCNA lower limit of detection (3%), thus defined as undetected by ichorCNA (ichorCNA-) (**Table S7**). As expected, we observed no statistically significant difference between ichorCNA TC estimations of ichorCNA-versus control samples (p=0.06, AUC=0.62) (**Fig. 2D**). The Proportion of Tumor-like Reads (PTR), a measure implemented in METER-detect to estimate the presence of tumor DNA signal (**Online Methods**), resulted in strongly statistically significant different levels in tumor versus control samples (p<1e-7, AUC=0.85) (**Fig. 2E)**. Using a Z-score statistics applied to the distribution of PTR in control samples, METER-detect using Z-score=3 classified 42% ichorCNA-samples, corresponding to 25 out of 60 samples, as METER+. Using a leave-one-out strategy applied to control samples, we evaluated FDR at different thresholds of Z-score, obtaining FDR<5% for Z-score=3 (**Fig. 2F)**. We then conducted extensive in-silico studies to investigate the performance of our method in relation to Z-score thresholds and coverage. Considering in-silico dilutions at TC=0, corresponding to 10 different pools of plasma samples from healthy donors, at each coverage level we observed a decrease in FDR with an increase in the Z-score threshold, as expected. Within the coverage range similar to this study (0.2-1X) and with Z=3, we observed an FDR of less than 10%, consistent with the leave-one-out study performed on real samples. The analysis also indicated a sensitivity of 100% and 60% for TC 1% and 0.5%, respectively. Notably, the sensitivity of METER-detect increased with the level of coverage, suggesting the possibility of detecting ctDNA at TC=0.1% with a sensitivity of 70% using Z=4 in WGBS 4x, while still maintaining an FDR of less than 10% (**Fig. 2G** and **Fig. S4)**.

### ER subtyping with METER-subtype

The possibility to analyze the ER status in the circulation of BC patients is emerging as a clinically relevant need. Indeed, ER status may change during metastatic tumor progression due to resistance to endocrine therapy and/or chemotherapy (*24*), and performing serial tissue biopsies is usually not feasible, especially in the case of brain metastases (*25*). We therefore hypothesized the existence of genome-wide patterns of DMR able to inform about the ER status, with the possibility to be exploited in ctDNA low-pass analysis. To do that, we implemented a computational strategy in METER, called METER-subtype, to classify each sample into ER+ or ER-. Our procedure involves the identification of subtype iDMR from the BASIS dataset (*18*), followed by the application of the Robust Partial Correlation (RPC) method as implemented in the EpiDISH tool (*26*) to deconvolute DNA-methylation signal from each lpWGBS samples into three components: ER+, ER- and normal cfDNA (**Online Methods**). Subtype iDMR showed distinctive patterns of DNA-methylation median levels of ER+ and ER-samples, as expected based on the procedure used for their selection (**Fig. 3A** and **Table S8**). Applying RPC to lpWGBS samples, we first observed that the ER+ and ER-component levels were consistent with the available immunohistochemistry (IHC) evaluation based on the most recent archived sample available (**Fig. 3B** and **Table S9**). The RPC component strongly associated with TC by METER-quant when expected IHC-based ER status was considered (R=0.9, p<1e-10 for both ER+ and ER-), while less evident for ER+ component in ER-samples (R=0.45, p=0.015) or no significant association for ER-component in ER+ samples (R=0, p=1) was obtained (**Fig. 3C**). Considering the difference between ER+ and ER-components in each sample, we observed evident differential distribution between ER+ and ER-components based on IHC status for samples classified with TC by METER-quant≥5% (n=53, p=0.004, AUC=0.86), while no difference was observed for METER-samples. Interestingly, our data suggest the possibility to infer ER status also in case of low TC values, as obtained for METER+ with TC<5% samples (p<0.05, AUC=0.73) (**Fig. 3D**). ER status was then assigned based on the predominant component between ER+ and ER-(METER-subtype) and compared to IHC. We observed that the accuracy of classification showed dependence on TC, with 88% (n=23) of misclassifications occurring for TC<5% (**Fig. S5**). Considering the 53 samples with METER-quant >5%, METER-subtype showed an accuracy of 0.94 (CI=0.83-0.99) in correctly classifying ER status (**Fig. 3E**), with a call rate of 0.92 due to 8% of cases (n=4) where RPC was unable to estimate ER+ or ER-component levels. Importantly, among the 8 patients with available BL/Prog samples and METER-quant>5% in both time points (all ER+ based on IHC), consistent subtypes were predicted at both time points (**Fig. S5**).

**Fig. 3.**
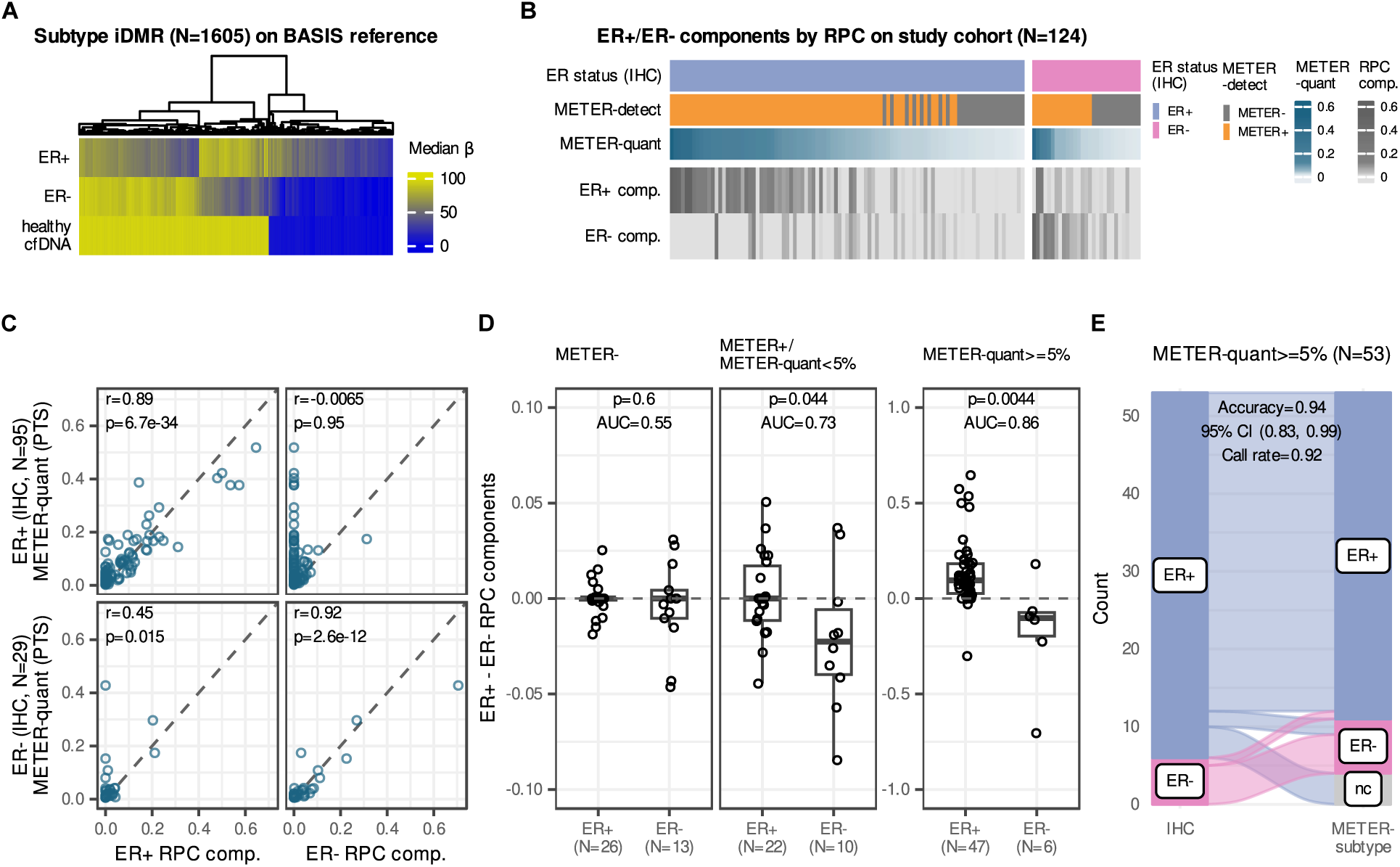
ER subtyping from cfDNA samples by METER-subtype. **(A)** Unsupervised clustering of selected subtype iDMR based on median β by DMR in reference ER+/HER2-(N=24) and ER-/HER2-(N=5) from BASIS dataset and healthy cfDNA samples (N=23) from Fox-Fisher et al. dataset. Clustering method: complete-linkage hierarchical clustering based on euclidean distance. **(B)** Heatmap showing ER+ and ER-components by RPC in the 124 mBC cfDNA samples within the study cohort. ER status by IHC, TC by METER-quant and classification by METER-detect are shown for each sample. **(C)** Scatter plots showing TC by METER-quant (y-axis) and ER+/ER-components by RPC (x-axis) for ER+ (top panels) and ER-(bottom panels) mBC samples based on IHC classification. R and p are from Pearson’s correlation test. **(D)** Boxplots showing the distribution of the difference between ER+ and ER-RPC components in ER+ and ER-BC samples, as classified by IHC, within the study cohort. Left panel: METER-samples (N=39); middle panel: METER+ samples with TC by METER-quant<5% (N=32); right panel: METER+ samples with TC by METER-quant>=5% (N=53). P is Wilcoxon Test p-value. **(E)** Alluvial plot showing concordance between ER subtyping by IHC and METER-subtype on samples with TC by METER-quant≥5% within the study cohort. Median β: median of beta by DMR (where beta by DMR is the mean of the beta values of CpGs within the specific DMR); RPC: robust partial correlation; IHC: immunohistochemistry; PTS: proportion of tumor-like sites; r: Pearson correlation coefficient; AUC: area under the curve; CI: confidence interval; mBC: metastatic breast cancer; TC: tumor content.

### Association of METER results with clinically relevant prognostic factors

In order to evaluate the potential utility of METER for clinical applications, we first investigated its results in association with clinical data collected from the patients included in this study. Using the number of metastatic sites at study entry as a proxy of tumor burden, we first observed that METER-quant at pre-treatment baseline positively correlated with this measure (Spearman’s rho=0.44, p<1e-3), and higher although not significant TC values were observed in patients with visceral disease vs bone or non-visceral (p=0.08) (**Fig. S6**). Interestingly, we did not observe a trend indicating an increase in TC for later treatment lines (**Fig. 4A**). All patients included in this study were previously characterized (*27–29*) for the number of CTC, for a total of 121 out of 124 samples. We observed a positive correlation between TC estimates by METER-quant and the number of CTC (rho=0.57, p<1e-11) (**Fig. 4B**). A significantly higher number of CTC was detected in METER-detect positive samples compared to negative samples (p<1e-6). In particular, only 8 out of 37 (22%) METER-detect negative samples showed CTC>0, with a range of 1-3. Importantly, significant association between the number of CTC and METER-detect classification was obtained when samples undetected by ichorCNA were considered (p=0.002) (**Fig. 4C**).

**Fig. 4.**
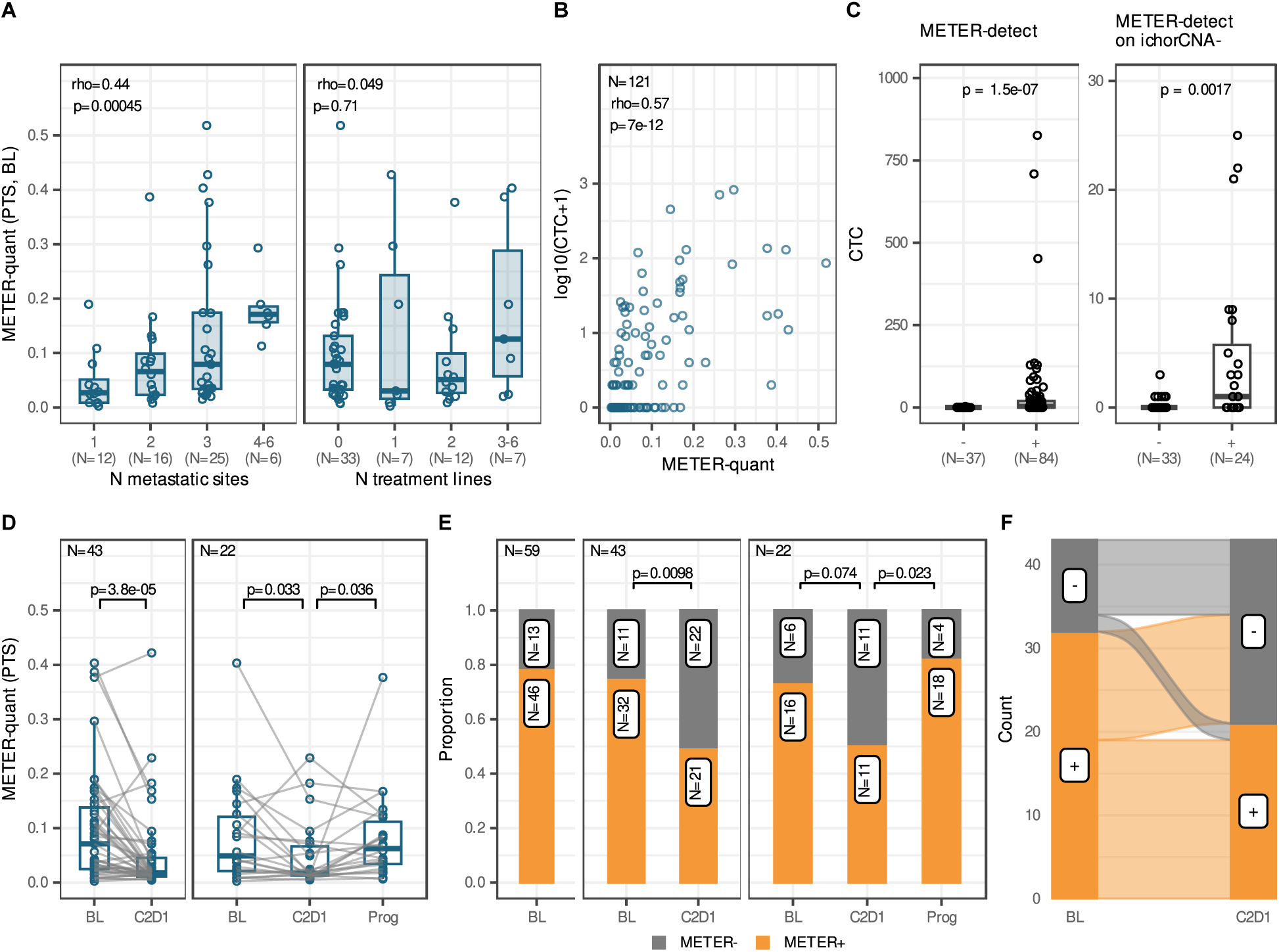
Association of METER estimates with clinically relevant prognostic factors. **(A)** Box plots showing the distribution of TC by METER-quant in mBC samples at BL (N=59) stratified by number of metastatic sites at study entry (left) and number of metastatic treatment lines at study entry (right). p is Pearson’s correlation Test p-value. **(B)** Scatterplot of TC by METER-quant (x-axis) versus the logarithm base 10 of the number of CTC in samples (y-axis). p is Pearson’s correlation Test p-value. **(C)** Box plots showing the distribution of the number of CTC in METER- and METER+ samples, for all samples (N=124, left panel) and for samples with TC by ichorCNA<3% (N=57, right panel) within the study cohort. p is Wilcoxon Test p-value. **(D)** Box plots showing the distribution of samples’ TC by METER-quant (y-axis) across clinical time points (x-axis) for patients having both BL and C2D1 time points (left panel, N=43) and for patients with complete longitudinal data (right panel, N=22) within the study cohort. p is paired Wilcoxon p-value. **(E)** Bar plots showing the proportion of METER+ cfDNA samples (y-axis) across clinical time points (x-axis) for all patients at BL (left panel, N=59), for patients having both BL and C2D1 time points (middle panel, N=43) and for patients with complete longitudinal data (right panel, N=22) within the study cohort. p is McNemar’s chi-squared test p-value. **(F)** Alluvial plot showing the dynamics of samples’ classification by METER-detect in patients having both BL and C2D1 time points (N=43) within the study cohort. Rho: Spearman’s correlation coefficient; TC: tumor content; CTC: circulating tumor cells; BL: pre-treatment baseline; C2D1: cycle 2 day 1; Prog: progression.

We then examined the capability of METER-quant for longitudinal monitoring of patients with mBC. In 43 patients having samples collected at both pre-treatment baseline (BL) and at day 1 of cycle 2 (C2D1), a statistically significant drop of TC by METER-quant at C2D1 versus BL was observed (paired p<1e-4), confirming expectedly sensitivity to treatment for the majority of the patients. Similar results were obtained using ichorCNA on the same samples (p<1e-3, **Fig. S7).** In 22 patients with complete longitudinal data, thus including the plasma sample collected at progression, significant differences of METER-quant for both C2D1 vs BL (p<1e-2) and sample collected at progression (Prog) vs C2D1 (p<0.05) were obtained (**Fig. 4D**). The tool ichorCNA showed a borderline significance for the comparison C2D1 vs BL (p=0.05), while no significant increase at Prog versus C2D1 (p=0.59) was obtained (**Fig. S7**). We then examined patients’ classification based on METER-detect across the plasma samples’ time points. We observed a significantly higher fraction of METER+ samples when samples were collected before starting a new line of treatment (74%, p=0.002, Fisher’s Exact test) and at progression (Prog, 82%, p=0.01) compared to C2D1 (49%) (**Fig. S6**). Results were confirmed in the 43 patients having both BL and C2D1 samples (p<0.01, McNemar test) and with complete longitudinal data (McNemar p=0.07 and p=0.02 for BL and Prog versus C2D1, respectively) (**Fig. 4E**). Comparable results were obtained using ichorCNA on all samples within the study cohort, while, considering matched samples only, significance was reached only in the 43 patients having both BL and C2D1 time points (**Fig. S7**). We then examined the dynamic change of ctDNA at C2D1 versus BL. Patients with available C2D1 were classified based on the detection of ctDNA provided by METER, as +/+ (if ctDNA was detected at both BL and C2D1), +/- (if ctDNA was detected at BL but not at C2D1), -/- (if ctDNA was not detected neither at BL nor at C2D1). We observed that the majority (81%, 9 out of 11) of patients classified as METER-at BL did not change their classification at C2D1, while 41% (13 out of 32) of METER+ patients at BL were classified as METER-at C2D1, consistent with previous findings (**Fig. 4F**). Similar results were obtained using ichorCNA classification (**Fig. S7**).

### ssociation of M T R with patients’ outcome

We then tested METER-detect in providing clinically meaningful stratification of patients by its association with the Progression Free Survival (PFS) and Overall Survival (OS) of patients included in this study. We therefore classified patients based on ctDNA detection at BL provided by METER-detect (METER+ or METER-) or ichorCNA (ichorCNA+ if TC by ichorCNA≥3%, ichorCNA-otherwise). Considering BL classification, METER+ patients showed a statistically significant shorter OS (Hazard Ratio (HR)=4.2, 95% Confidence Interval [CI]=2-9, p<0.0001) than METER-, and demonstrated superior prognostic ability than ichorCNA classification (HR=2.1, CI=1.3-3.8, p=0.002). Considering the ichorCNA-patients (n=23), despite the small number a trend for METER+ towards worse OS was observed (HR=2.2, CI=0.9-5.5, p=0.09) (**Fig. 5A**). Examining the association with PFS, a worse outcome for METER+ patients, with a HR of 3.8 (CI=1.8-8.0, p<0.001), was observed. In contrast, no significant association was obtained using stratification provided by ichorCNA. Notably, for ichorCNA-patients a significant worse PFS for patients classified as METER+ compared to METER-was still obtained (HR=3.9, CI=1.5-10.0, p=0.003) (**Fig. 5B**). We then investigated the dynamic change of ctDNA considering the classification provided by METER and ichorCNA at C2D1 versus BL. Patients -/+, for whom ctDNA was not detected at BL, but detected at C2D1, were excluded because of the small number of observations (n=3 in total, n=1 for both METER-detect and ichorCNA). Considering OS, we observed superior ability of METER-detect (p<0.001) in stratifying patients compared to ichorCNA (p=0.02), especially due to the more reliable identification of patients with absent ctDNA (median OS 5.9mo and 2.4mo for METER-detect and ichorCNA, respectively). Notably, considering the 15 patients classified as ichorCNA -/-, METER-detect was still able to provide statistically significant stratification (p=0.01) (**Fig. 5C**). Similar results were obtained considering the PFS as outcome, where we confirmed the superior ability of METER-detect classification (p<0.001) in stratifying patients compared to ichorCNA (p=0.002) and statistically significant stratification also for the more challenging ichorCNA -/- patients (p=0.003) (**Fig. 5D**). Notably, in multivariate analyses controlling for relevant covariates, including BC subtype, number of previous treatment lines at study entry, number of metastatic sites at study entry and type of metastatic disease, METER-detect at T0 remained significantly associated with both PFS (p=0.006) and OS (p=0.002), and the same when considering BL/C2D1 dynamics (**Table S10**).

**Fig. 5.**
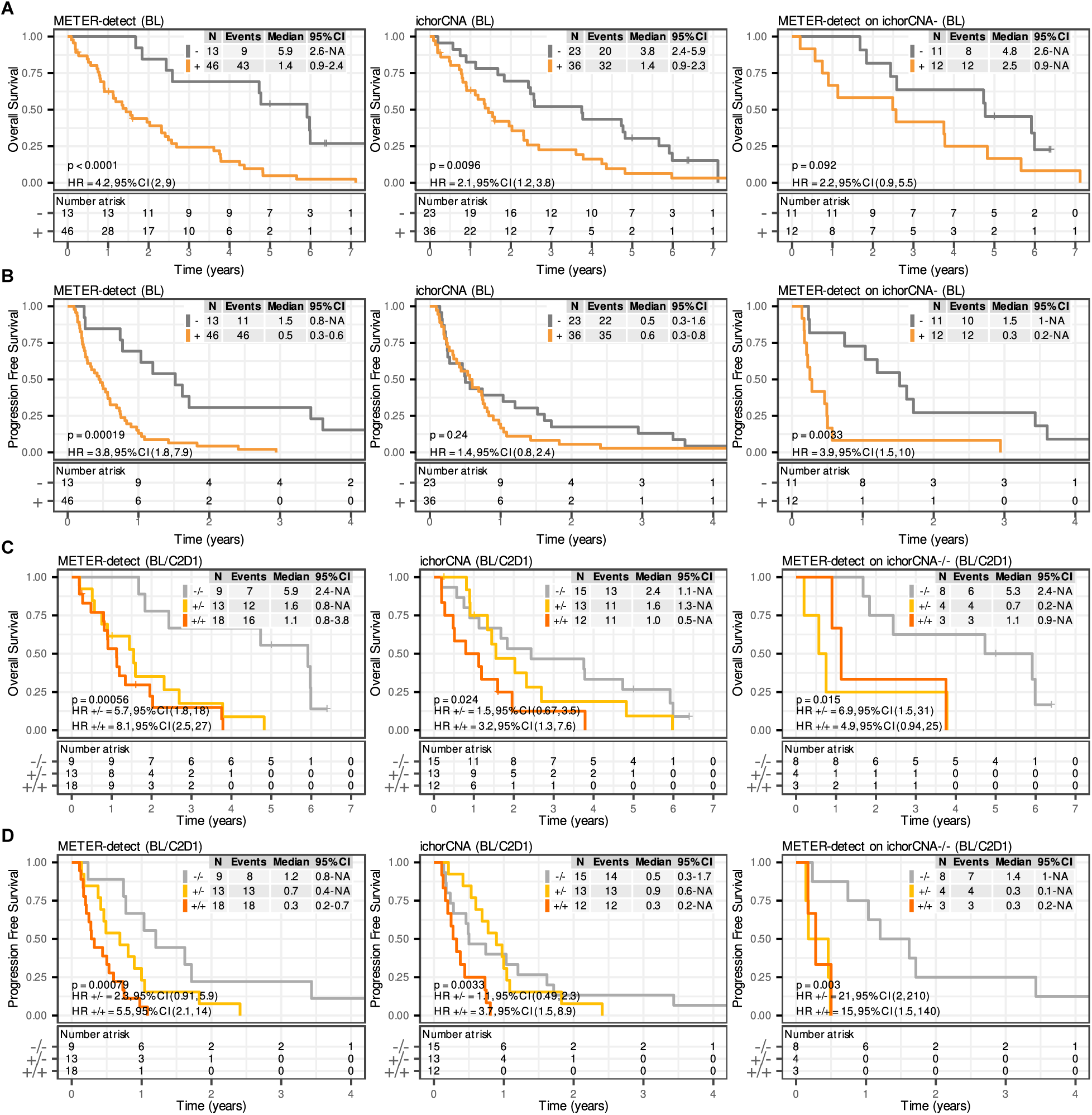
Association of METER-detect classification with patients’ outcome. Kaplan-Meier curves showing OS **(A)** and PFS **(B)** of patients stratified according to samples’ classification at BL by METER-detect (METER+ or -) and ichorCNA (ichorCNA+ or -), for all patients (N=59, left and middle panels respectively), and for ichorCNA-patients at BL (N=23, right panel) within the study cohort. Kaplan-Meier curves showing **(C)** OS and **(D)** PFS of patients stratified according to samples’ classification at BL and C2D1 time points (BL/C2D1) by METER-detect (METER+/+, +/- or -/-) and ichorCNA (ichorCNA +/+, +/- or -/-), for all patients having both BL and C2D1 time points (N=43, left and middle panels respectively), and for ichorCNA-/- patients at BL/C2D1 (N=15, right panel) within the study cohort. BL: pre-treatment baseline; C2D1: cycle 2 day 1; p: Log Rank Test p-value; CI: confidence interval; HR: hazard ratio.

## Discussion

In this study, we proposed a new strategy combining the use of low-pass methylome sequencing followed by a novel computational tool named METER to perform comprehensive analysis of TC, including quantification, detection and transcriptional activity, such as molecular subtyping. We showed the utility of our approach applying METER to lpWGBS data (0.8X) of plasma samples from 124 patients with mBC and 30 healthy donors. Our approach relies on the characteristics of DNA-methylation being prevalent, pervasive and cell-type-specific in cancer cells (*30*). These distinctive features enable genome-wide (“horizontal”) analysis of sequencing data, in contrast to the majority of ctDNA strategies relying on depth of coverage of specific sites and/or regions (“vertical”) to draw any inference. This type of analysis makes our strategy conceptually similar to ichorCNA but with the advantage of being applicable also in case of genomes characterized by low-to-absent CN burden. In addition, as reported in our feasibility study (**Supplementary Text**), lpWGBS can be used to obtain accurate estimation of CNA profiling, with the possibility to exploit multi-modal analysis of these data, consistent with previous work exploiting both CN and DNA-methylation alterations for tumor sample analysis (*31, 32*). The use of lpWGBS coupled with METER has several practical advantages compared to other approaches proposed for ctDNA analysis. It is reproducible requiring only bisulfite treatment of isolated DNA followed by NGS library preparation and sequencing. It requires very low amount of input DNA (10-20ng), typically manageable from 1mL of plasma. Compared to other methodologies implemented by currently available commercial kits is economically convenient. Our approach is tumor tissue-agnostic, not depending on the analysis of tumor tissue-specific (e.g. hotspot) alterations, but exclusively on pre-defined iDMR and iDMS. Further, the analysis of different sets of specific iDMR and iDMS (through the analysis of publicly available datasets) allows for its potential application to different tumor types or subtypes. Another advantage of our approach is that the computational strategies employed in METER do not require model training on lpWGBS data, except for estimating the mean and variance on the control samples used in the METER-detect module (mean and variance precomputed values are also provided within the METER tool). In contrast, many of the other epigenetics-based approaches for ctDNA analysis depend on the training of machine learning classifiers, including methods analysing nucleosome positioning (*16*), fragmentomics (*12*), or MeDIP profiling (*11*).

We demonstrated that TC estimations by METER-quant were consistent with those provided by the state-of-the-art tool ichorCNA, and using ad-hoc artificial samples we observed reliable TC estimations below ichorCNA’s lower limit of detection, with linearity maintained down to 0.5-1%. METER-detect applied to ichorCNA undetected samples identified tumor signal in almost half of them, yet maintaining FDR <5%. Through a comprehensive in-silico study, for coverage levels similar to our dataset we estimated a lower limit of detection for METER-detect at approximately 1%, with promising performance for higher coverage levels. In fact, for coverage in the range of 4x, a sensitivity exceeding 90% for TC=0.1% and an FDR below 10% were estimated. Despite the increase in coverage and thus higher costs related to sequencing and data, this result represents a significant improvement over ichorCNA, where the authors reported a sensitivity of 95% and an FDR of 9% for TC 3%, thus potentially enabling the detection of TC levels one magnitude lower.

Exploiting DNA-methylation regions as a proxy of specific transcriptional programs, we here demonstrated the application of METER-subtype to infer the ER status in circulation accurately. In samples with METER-quant>5%, METER-subtype inferred the ER status in circulation with 94% accuracy, abstaining on 8% (n=4) of samples. We observed misclassifications in three patients. One patient had an estimated TC close to 5%, while the other two had notably higher TC. Specifically, of those with higher TC, one patient (MS10) identified as ER+ by IHC was predicted as ER-, while the other patient (MS43) classified as ER-by IHC was predicted as ER+. Interestingly, differently from the majority of cases both ER+ and ER-components were detected in these samples (MS10: ER+=0.01, ER-=0.31 and MS43: ER+=0.21, ER-=0.03) (**Fig. S5**) and both patients had advanced disease involving multiple metastatic sites (3 and 4 metastatic sites for MS43 and MS10 respectively). The accuracy of ER status provided by METER-subtype was in line with the tool Griffin working on nucleosome positioning (*16*). However, our method is based on the deconvolution of DNA-methylation signal of predefined specific DMR and doesn’t require the training of a machine learning classifier to infer subtype. Additionally, we observed acceptable performance for METER+ samples with TC<5%, thereby demonstrating subtyping in circulation in an extremely challenging range of TC levels for this task. It is also important to note that our dataset mostly includes patients with early metastatic disease. Considering the evaluable samples in our dataset (METER-quant ≥ 5%), the median TC was 13% (IQR=8-18%). Conversely, in the Griffin dataset the TC for evaluable (ichorCNA>5%) samples was significantly higher (p=0.002, IQR=10-39%) (**Fig. S8**). As the accuracy in ER subtyping depends on TC, we expect promising applications of METER-subtype for the non-invasive monitoring of ER status in later lines of treatments, where more frequent discordances in ER status may be expected due to acquired resistance mechanisms caused by longer exposure to endocrine therapy and/or chemotherapym (*33, 34*) and to detect subtype switching (*35, 36*), especially those occurring in patients with brain metastases (*37*) . Recently, the possibility to infer transcriptional programs with clinical relevance from DNA features estimated from low-pass WGS was shown, including ER status and RB-loss (*38*). However, the authors reported failure rate up to 20% for the method in case of TC in the range of 5-10% and lower accuracy (AUC=0.8) compared to this and previous study (*16*) for inferring ER status.

Most studies reporting new methodologies for ctDNA analysis typically focus on demonstrating advantages in terms of technical performance, with a lack of evaluation regarding their relevance for clinical applications. In this study we demonstrated the clinical relevance of METER’s prediction by examining association with metastatic burden, CTC and its utility for longitudinal monitoring. In addition, METER-detect offered clinically meaningful and superior stratification of patients in terms of OS and PFS, considering both the BL time point and the dynamic change at C2D1 compared to BL. Of note, METER-detect provided refined stratification for patients whose plasma samples were undetected by ichorCNA, thus indicating the reliability of our approach for potential clinical applications. Recent data have been reported on the use of TC≥10% as a prognostic independent factor in advanced colorectal, prostate, non-small cell lung and breast cancer (*5*). In our cohort, we confirmed TC detection by METER-detect as independent prognostic factor. Specifically, this result obtained for METER-detect classification suggests that a markedly lower threshold of TC (1% from our in-silico study) than previously reported (10% from the Reichert et al. study (*5*)) should be consider informative for patients’ stratification, in line with other studies (*13*).

Our analyses are dependent on a catalog of DMR identified from WGBS, which is currently limited in availability to the scientific community. In fact, the vast majority of cancer DNA-methylation data has indeed been generated using array data. As reported in **Supplementary Text** and **Fig. S9** and **S10**, using DMR detected by Rocker-meth (*39*) in TCGA-BRCA dataset (*30*) we observed consistent findings for both METER-quant and METER-detect. The association of METER-detect exploiting these TCGA-BRCA derived DMR with clinical outcomes resembles those obtained using WGBS. Although overall consistent trends Although consistent trends were observed overall, using iDMR identified through array data lower performance in patient stratification was obtained for ichorCNA-patients when compared to sequencing-based iDMR (**Fig. S11**), likely due to the lower quantity and reliability of array-based iDMR. Despite this, we anticipate no significant limitations that would hinder the application of our approach for the analysis of samples from other tumor types.

In early settings, typically lower TC is detected posing a limitation for the use of lpWGBS for applications such as minimal residual disease (MRD) or monitoring disease during adjuvant treatment. However, it’s important to note that commonly used strategies in these contexts include tumor tissue informed analyses, high coverage sequencing of large DNA and/or DNA-methylation sites or WGS (*40–42*) . With obvious limitations due to in-silico modeling, our synthetic study suggests potential improvement of sensitivity with increasing coverage, with the advantage of our approach being tumor tissue agnostic and not assisted by machine learning classifiers, both features largely desirable for liquid biopsy applications in this setting.

To conclude, our strategy represents a new highly reproducible and cost-effective approach that could facilitate the analysis of TC in plasma samples from patients with metastatic cancer, as well as providing information about relevant transcriptional programs, and enabling early identification of patients with the higher risk of resistance to metastatic treatment.

## Material and Methods

### Plasma samples collection and cfDNA extraction

A total of 30 healthy donors and 59 patients with metastatic breast cancer (mBC) were analysed. For mBC patients, a total of 124 samples were collected at distinct time points: baseline pre-treatment (BL, n = 59), day1 of the second cycle of treatment (C2D1, n= 43), and at disease progression (Prog, n= 22). cfDNA was extract by QIAmp Circulating Nucleic acid Kit (Qiagen) and quantified by Qubit (Qubit dsDNA High Sensitivity Kit, Life Technologies). Extracted cfDNA samples were stored at -80°C until the bisulfite conversion. Approximately 20 ng (mean: 19.8 ng; range: 10.52 ng - 22.9 ng) of cfDNA was bisulfite converted using the EZ DNA Methylation-Lightning Kit (Zymo Research) according to the manufacturer’s instructions. Samples with volume exceeded the 20ul input for conversion was concentrated by SpeedVac DNA130 (Thermo Fisher Scientific) at 35°C. Bisulfite converted cfDNA was eluted into 15ul of Low EDTA TE and used for library preparation. Libraries were prepared using the Accel-NGS Methyl-Seq DNA Library Kit (Swift Biosciences, 30096), the Swift Normalase Unique Dual Indexing Primer Plates (Swift Biosciences, X91384-PLATE) and the KAPA HiFi HotStart Uracil (Roche) according to the manufacturer’s instructions. 13 cycles of indexing PCR were conducted in order to achieve at least 300 ng of product. Amplified libraries were purified using 0.75x volume of SPRIselect beads (Beckman Coulter). Libraries’ size distribution and concentration were analyzed respectively by Bioanalyzer DNA High Sensitivity Kit (Agilent) and Qubit (Qubit dsDNA High Sensitivity Kit, Life Technologies) before pooling and sequencing. Sequencing was performed with 15-25% PhiX (Illumina) spike in at 2×150 bp on Illumina Novaseq 6000.

### *In-vitro* and *in-silico* artificial samples

In vitro artificial dilutions were generated mixing DNA extracted from luminal T47D BC cell line culture medium, as a surrogate of circulating tumor DNA (s-ctDNA), and a human mixed genomic DNA (PROMEGA) (0, 0.1, 0.5, 1, 2.5 and 5%). s-ctDNA was extracted by QIAmp Circulating Nucleic acid Kit (Qiagen) and quantified by Qubit dsDNA High Sensitivity Kit, (Life Technologies).

*In-silico* artificial dilutions were generated by combining sequenced reads from WGBS of the luminal MCF7 breast cancer cell line, downloaded from Sequence Read Archive (SRA, GEO Accession: GSM3336908), with sequenced reads from pooled lpWGBS of 20 controls from the MIMESIS lpWGBS dataset. Reads from these two sources were randomly sampled and mixed in proportions to simulate increasing TC and sequence coverage, using SAMtools version 1.16.1 (*43*). Specifically, TC values ranging from 0 to 0.02 were simulated, generating 10 dilutions for each TC point, within distinct simulated coverage levels: 0.7X, 1.3X, 2.7X, and 4.0X. lpWGBS data from 20 out of 30 controls available were used to generate the dilutions, while data from the remaining 10 controls were pooled and subsequently used to generate 10 additional artificial controls, for each simulated coverage interval, through the random sampling of reads using SAMtools.

### Processing of cfDNA-methylation data

Sequenced reads were processed through the ’methylseq’ pipeline version 1.6.1 within the nf-core project version 2.2 (*44*), using the Bismark workflow (*45*),. Briefly, reads were aligned to the hg38 genome version using Bismark (*45*), trimmed using accel-ngs methyl-seq specific parameters by Trim Galore (https://www.bioinformatics.babraham.ac.uk/projects/trim_galore/). Deduplication and methylation calls were obtained using Bismark. For in silico artificial dilutions, only the final step of methylation calls using Bismark was applied, as the sequenced reads used in creating the dilutions had already undergone processing. Reference TC estimate was determined applying ichorCNA (*9*) to lpWGBS data, using the following parameters: mapping quality=20, window size=500 Kb, allowing normal fraction up to 0.99 and using the panel built on the lpWGBS data from the 30 healthy cfDNA samples from the MIMESIS lpWGBS dataset.

### Selection of iDMS and iDMR

To define BC-specific Differentially Methylated Sites (DMS) and Regions (DMR), Rocker-meth (*39*) was applied to beta from WGBS profiles of 30 BC tissue (*18*) (BASIS dataset) and 23 healthy cfDNA (*19*). Subsequently, a series of filtering criteria were applied in order to select TC informative DMS (iDMS) and DMR (iDMR).

For iDMS, the following criteria were used:

- Area under curve (AUC) of a receiver operating characteristic (ROC) curve, separating tumor from control samples, higher than 0.80 (hyper-methylated sites) or lower than 0.2 (hypo-methylated sites);
- Difference of mean beta values between tumor and control samples greater than 0.4 (where beta value is defined as the per-site proportion of methylated signal over the total methylation signal;
- Third quartile of beta values in control samples lower than 0.1 (hyper-methylated) or first quartile of beta values in control samples above 0.9 (hypo-methylated);

leading to the selection of 45K hyper and 165K hypo iDMS (**Table S2**). For iDMR, the following criteria were used:

- FDR<0.05;
- difference between mean of beta by DMR values between tumor and control samples greater than 20 (where beta by DMR values are computed as the mean beta values of CpG sites within the specific DMR;
- third quartile of beta by DMR values in control samples lower than 0.1 (hyper-methylated) or first quartile of beta by DMR values in control samples above 0.9 (hypo-methylated);

leading to a selection of 1,778 hyper and 6,703 hypo iDMR (**Table S6**).

### The METER computational tool

METER is a computational tool to analyze TC exploiting iDMS and iDMR in lpWGBS data of cfDNA samples. It comprises three modules: 1) METER-quant, to measure circulating tumor DNA (ctDNA), based on tumor-specific iDMS; 2) METER-detect, to classify samples as ctDNA+ or ctDNA-(that is ctDNA is detected or not), based on tumor-specific iDMR; 3) METER-subtype to infer specific subtype from ctDNA, based on tumor subtype-specific iDMR.

#### METER-quant

In this module, the proportion of tumor-like sites (PTS) by sample is measured from lpWGBS as a proxy of sample’s TC. PTS for each sample is computed as the ratio between fully methylated (beta=100%) hypermethylated or fully unmethylated (beta=0%) hypomethylated iDMS (that is iDMS supporting a tumor-like methylation signal) to the total fully methylated/unmethylated iDMS covered. To minimize the effect of bisulfite conversion errors and increase signal specificity, only reads with alpha value of 100%, that is reads showing only methylated or unmethylated CpG sites, and covering a minimum of 6 CpG sites (user-selectable parameter) are considered for the computation of beta values.

In addition to the direct estimate of TC through PTS measurement, a linear model was trained on 94 (60%) randomly selected samples (N=76 tumors an N=18 controls) from the study cohort, using PTS measure as predictor and 0 TC or TC estimated by ichorCNA as reference, for control and patients cfDNA samples respectively. The model was then tested on the remaining 60 (40%) samples (N=48 tumors and N=12 controls). Predicted negative values were set to TC=0%. Parameters and results are shown in **Fig. S3** and **Table S5.**

#### METER-detect

Within this module, the proportion of tumor-like sequenced reads (PTR) is computed from lpWGBS data for each sample, and a z-score approach, based on the distribution of this measure in control samples (reference model), is then used to classify samples as either ctDNA positive (METER+) or ctDNA negative (METER-). As in METER-quant module, to minimize the effect of bisulfite conversion errors and increase signal specificity only reads with alpha value of 100% and covering a minimum of 6 CpG sites (user-selectable parameter) are considered for PTR computation. Specifically, PTR for each sample is computed as the proportion of fully methylated reads within selected hyper-iDMR and fully unmethylated reads within selected hypo-iDMR (that is reads supporting a tumor-like methylation signal) over the total reads with alpha=100% within selected iDMR. Subsequently, the Z-score is computed with respect to control samples, and a sample is classified as ctDNA-positive when its Z-score exceeds a threshold (user-selectable parameter).

Prior PTR computation, an additional filtering to iDMR is applied, excluding iDMR that exhibit unexpected methylation signal in control samples. Specifically, any hyper-iDMR covered by one or more fully methylated reads (with a minimum of 6 CpGs), exhibiting an unexpected tumor-like methylation pattern, in at least three control samples are excluded. Likewise, any hypo-iDMR covered by one or more fully unmethylated reads (with a minimum of 6 CpGs) in at least three control samples are also excluded. For the 30 samples considered in this study, a total of 40 iDMR were excluded (**Table S11**).

#### METER-subtype

To assess the Estrogen Receptor (ER) status of cfDNA samples, we applied Robust Partial Correlation (RPC) method to the subset of subtype-specific iDMR. Using WGBS samples from the BASIS dataset, subtype-specific iDMR were defined as those ones showing a mean beta difference between ER+/HER2- and ER−/HER2-samples greater than 20 (**Table S8**). A reference model including three components (ER+, ER- and healthy cfDNA) was built by computing the median of the beta by DMR values (where beta by DMR values are computed as the mean beta values of CpG sites within the specific DMR) in ER+/HER2- and ER-/HER2- samples from the BASIS dataset, and in healthy cfDNA samples from Fox-Fisher et al. dataset. For lpWGBS data a beta table was generated, comprising beta by DMR values of subtype iDMR for each sample. In this case beta by DMR values were computed by treating all CpGs within a specific DMR collectively as a single CpG. Therefore, the beta value for each DMR was computed as the ratio of methylated signals to the total methylation signals for all CpGs within that DMR. Finally, the deconvolution method RPC implemented in the EpiDISH R package (*26*) was used to estimate the proportion of each component in each sample. For each sample, ER status classification was performed based on the maximum component between ER+ and ER- estimated by RPC.

### Analysis of False Discovery Rate in METER-detect

The False Discovery Rate (FDR) of METER-detect was assessed through a leave-one-out strategy applied to PTR in control samples. At each iteration, one control sample was classified as METER+ or METER-based on the reference model (mean and sd for Z-score and iDMR cleaning) estimated using the other 29 samples. The relationship between FDR and Z-score thresholds, from 1 to 6, was investigated. Following this procedure, we classified 0 out of 30 controls as ctDNA+ using a Z-score=3, supporting an FDR of less than or equal to 0.05 in our dataset.

### Evaluation of coverage levels on METER-detect sensitivity

To evaluate the impact of coverage on detection sensitivity and FDR, in silico artificial dilutions at increasing coverage levels were generated (see section *In-vitro* and *in-silico* artificial samples). To account for the reduced variability of DNA methylation signals in these artificial control samples compared to real healthy cfDNA samples, we estimated the mean and standard deviation for calculating Z-scores at each coverage level using the 10 artificial controls that were not included in generating the dilutions.

### Statistical Analysis

Statistical analyses were performed with the R software environment for statistical computing and graphics (R core team, 2022 https://www.R-project.org/). Survival analyses used the Cox proportional hazard method, with the logrank test. OS was computed from the date of informed consent to death from any cause, PFS was computed from informed consent to disease progression or death. A multivariate Cox regression model was fitted to evaluate the independent effect of each covariate on OS and PFS. Relevant covariates were selected according to clinical relevance. These include: the number of treatment lines received for metastatic disease (no previous treatment vs 1 or more lines) and the number of metastatic anatomic sites (single site vs multiple sites). Mann-Whitney (for 2 groups) or Kruskal-Wallis (for 3 groups or more) tests were used to assess the significance of the relationship between continuous and categorical variables. All correlations are Spearman’s correlations. Significances between two categorical variables were assessed using the Fisher’s exact test.

## Supporting information

Supplementary Text and Figures to METER manuscript

## Data availability

Processed data of cfDNA lpWGBS samples and the code to reproduce the analyses presented in the manuscript are available at https://doi.org/10.5281/zenodo.10964554. METER was developed as a package for R and is available under MIT license at https://github.com/caos-lab-unifi/METER.

## Acknowledgments

The authors thank the members of the bioinformatics unit and the translational research unit of the Hospital of Prato (Prato) and the laboratory of computational and functional oncology (University of Trento) for fruitful discussions. The authors thank Roberto Bertorelli and Veronica De Sanctis from the NGS core facility of CIBIO (University of Trento) for their precious help on setting up the sequencing experiments. The authors thank all the patients who participated in the program and their families.

## Ethics declarations

The trials included in this study received approval from the local institutional ethics committee of the Hospital of Prato and by the Area Vasta Toscana Centro local Ethics Committee (CEAVC Em.2021-CEAVC study 15108), in accordance with Helsinki Declaration. Written informed consent was prospectively obtained from all patients participating in these trials.

## Authors’ contributions

M.P. developed METER and conducted bioinformatic analysis, under the supervision of M.B. and F.D. M.B. conceived the study and the main conceptual ideas and supervised the study with F.D. F.G. and A.N. conducted all molecular experiments, including samples’ processing, setup and execution of the low-pass methylomes. F.G. conducted CTC analyses. I.M. and L.M. supervised molecular analyses. S.D.D., L.L., M.P., G.S., E.R., L.M. and L.B contributed to patients’ enrollment, collection of clinical data and performed clinical investigation. D.R., C.B. G.M.F. performed bioinformatic and statistical analyses. M.B., F.D. and L.B. obtained project fundings. M.B., M.P., F.D. wrote the original draft, all authors revised and edited the manuscript. All authors have read and approved the manuscript.

## References

1. Cescon, D. W., Bratman, S. V., Chan, S. M. & Siu, L. L. Circulating tumor DNA and liquid biopsy in oncology. Nat. Cancer 1, 276–290 (2020).

2. Siravegna, G., Marsoni, S., Siena, S. & Bardelli, A. Integrating liquid biopsies into the management of cancer. Nat. Rev. Clin. Oncol. 14, 531–548 (2017).

3. Gouda MA, Janku F, Wahida A, Buschhorn L, Schneeweiss A, Abdel Karim N, De Miguel Perez D, Del Re M, Russo A, Curigliano G, Rolfo C, Subbiah V. Liquid Biopsy Response Evaluation Criteria in Solid Tumors (LB-RECIST). Ann. Oncol. Off. J. Eur. Soc. Med. Oncol. 35, 267–275 (2024).

4. Ignatiadis, M., Sledge, G. W. & Jeffrey, S. S. Liquid biopsy enters the clinic - implementation issues and future challenges. Nat. Rev. Clin. Oncol. 18, 297–312 (2021).

5. Reichert ZR, Morgan TM, Li G, Castellanos E, Snow T, Dall’Olio FG, Madison RW, Fine AD, Oxnard GR, Graf RP, Stover DG. Prognostic value of plasma circulating tumor DNA fraction across four common cancer types: a real-world outcomes study. Ann. Oncol. (2022) doi:10.1016/j.annonc.2022.09.163.

6. Stover DG, Parsons HA, Ha G, Freeman SS, Barry WT, Guo H, Choudhury AD, Gydush G, Reed SC, Rhoades J, Rotem D, Hughes ME, Dillon DA, Partridge AH, Wagle N, Krop IE, Getz G, Golub TR, Love JC, Winer EP, Tolaney SM, Lin NU, Adalsteinsson VA. Association of Cell-Free DNA Tumor Fraction and Somatic Copy Number Alterations With Survival in Metastatic Triple-Negative Breast Cancer. J. Clin. Oncol. Off. J. Am. Soc. Clin. Oncol. 36, 543–553 (2018).

7. Romanel A, Gasi Tandefelt D, Conteduca V, Jayaram A, Casiraghi N, Wetterskog D, Salvi S, Amadori D, Zafeiriou Z, Rescigno P, Bianchini D, Gurioli G, Casadio V, Carreira S, Goodall J, Wingate A, Ferraldeschi R, Tunariu N, Flohr P, De Giorgi U, de Bono JS, Demichelis F, Attard G. Plasma AR and abiraterone-resistant prostate cancer. Sci. Transl. Med. 7, 312re10–312re10 (2015).

8. Orlando F, Romanel A, Trujillo B, Sigouros M, Wetterskog D, Quaini O, Leone G, Xiang JZ, Wingate A, Tagawa S, Jayaram A, Linch M; PEACE Consortium; Jamal-Hanjani M, Swanton C, Rubin MA, Wyatt AW, Beltran H, Attard G, Demichelis F. Allele-informed copy number evaluation of plasma DNA samples from metastatic prostate cancer patients: the PCF_SELECT consortium assay. NAR Cancer 4, zcac016 (2022).

9. Adalsteinsson VA, Ha G, Freeman SS, Choudhury AD, Stover DG, Parsons HA, Gydush G, Reed SC, Rotem D, Rhoades J, Loginov D, Livitz D, Rosebrock D, Leshchiner I, Kim J, Stewart C, Rosenberg M, Francis JM, Zhang CZ, Cohen O, Oh C, Ding H, Polak P, Lloyd M, Mahmud S, Helvie K, Merrill MS, Santiago RA, O’Connor EP, Jeong SH, Leeson R, Barry RM, Kramkowski JF, Zhang Z, Polacek L, Lohr JG, Schleicher M, Lipscomb E, Saltzman A, Oliver NM, Marini L, Waks AG, Harshman LC, Tolaney SM, Van Allen EM, Winer EP, Lin NU, Nakabayashi M, Taplin ME, Johannessen CM, Garraway LA, Golub TR, Boehm JS, Wagle N, Getz G, Love JC, Meyerson M. Scalable whole-exome sequencing of cell-free DNA reveals high concordance with metastatic tumors. Nat. Commun. 8, 1324 (2017).

10. Diaz, L. A. & Bardelli, A. Liquid biopsies: genotyping circulating tumor DNA. J. Clin. Oncol. Off. J. Am. Soc. Clin. Oncol. 32, 579–86 (2014).

11. Shen SY, Singhania R, Fehringer G, Chakravarthy A, Roehrl MHA, Chadwick D, Zuzarte PC, Borgida A, Wang TT, Li T, Kis O, Zhao Z, Spreafico A, Medina TDS, Wang Y, Roulois D, Ettayebi I, Chen Z, Chow S, Murphy T, Arruda A, O’Kane GM, Liu J, Mansour M, McPherson JD, O’Brien C, Leighl N, Bedard PL, Fleshner N, Liu G, Minden MD, Gallinger S, Goldenberg A, Pugh TJ, Hoffman MM, Bratman SV, Hung RJ, De Carvalho DD.. Sensitive tumour detection and classification using plasma cell-free DNA methylomes. Nature 563, 579–583 (2018).

12. Cristiano S, Leal A, Phallen J, Fiksel J, Adleff V, Bruhm DC, Jensen SØ, Medina JE, Hruban C, White JR, Palsgrove DN, Niknafs N, Anagnostou V, Forde P, Naidoo J, Marrone K, Brahmer J, Woodward BD, Husain H, van Rooijen KL, Ørntoft MW, Madsen AH, van de Velde CJH, Verheij M, Cats A, Punt CJA, Vink GR, van Grieken NCT, Koopman M, Fijneman RJA, Johansen JS, Nielsen HJ, Meijer GA, Andersen CL, Scharpf RB, Velculescu VE. Genome-wide cell-free DNA fragmentation in patients with cancer. Nature 570, 385–389 (2019).

13. Giampaolo Bianchini; Luca Malorni; Grazia Arpino; Alberto Zambelli; Fabio Puglisi; Lucia Del Mastro; Marco Colleoni; Filippo Montemurro; Giulia Bianchi; Ida Paris; Giacomo Allegrini; Marina Elena Cazzaniga; Michele Orditura; Claudio Zamagni; Stefano Tamberi; Daniela Castelletti; Matteo Benelli; Maurizio Callari; Angela Santoro; Michelino De Laurentiis. Abstract GS3-07: Circulating tumor DNA (ctDNA) dynamics in patients with hormone receptor positive (HR+)/HER2 negative (HER2-) advanced breast cancer (aBC) treated in first line with ribociclib (R) and letrozole (L) in the BioItaLEE trial. Cancer Res. 82, GS3-07-GS3-07 (2022).

14. Baca SC, Seo JH, Davidsohn MP, Fortunato B, Semaan K, Sotudian S, Lakshminarayanan G, Diossy M, Qiu X, El Zarif T, Savignano H, Canniff J, Madueke I, Saliby RM, Zhang Z, Li R, Jiang Y, Taing L, Awad M, Chau CH, DeCaprio JA, Figg WD, Greten TF, Hata AN, Hodi FS, Hughes ME, Ligon KL, Lin N, Ng K, Oser MG, Meador C, Parsons HA, Pomerantz MM, Rajan A, Ritz J, Thakuria M, Tolaney SM, Wen PY, Long H, Berchuck JE, Szallasi Z, Choueiri TK, Freedman ML. Liquid biopsy epigenomic profiling for cancer subtyping. Nat. Med. 29, 2737–2741 (2023).

15. Franceschini GM, Quaini O, Mizuno K, Orlando F, Ciani Y, Ku SY, Sigouros M, Rothmann E, Alonso A, Benelli M, Nardella C, Auh J, Freeman D, Hanratty B, Adil M, Elemento O, Tagawa ST, Feng FY, Caffo O, Buttigliero C, Basso U, Nelson PS, Corey E, Haffner MC, Attard G, Aparicio A, Demichelis F, Beltran H. Noninvasive Detection of Neuroendocrine Prostate Cancer through Targeted Cell-free DNA Methylation. Cancer Discov. 14, 424–445 (2024).

16. Doebley AL, Ko M, Liao H, Cruikshank AE, Santos K, Kikawa C, Hiatt JB, Patton RD, De Sarkar N, Collier KA, Hoge ACH, Chen K, Zimmer A, Weber ZT, Adil M, Reichel JB, Polak P, Adalsteinsson VA, Nelson PS, MacPherson D, Parsons HA, Stover DG, Ha G. A framework for clinical cancer subtyping from nucleosome profiling of cell-free DNA. Nat. Commun. 13, 7475 (2022).

17. Romagnoli D, Nardone A, Galardi F, Paoli M, De Luca F, Biagioni C, Franceschini GM, Pestrin M, Sanna G, Moretti E, Demichelis F, Migliaccio I, Biganzoli L, Malorni L, Benelli M. MIMESIS: minimal DNA-methylation signatures to quantify and classify tumor signals in tissue and cell-free DNA samples. Brief. Bioinform. (2023) doi:10.1093/bib/bbad015.

18. Brinkman AB, Nik-Zainal S, Simmer F, Rodríguez-González FG, Smid M, Alexandrov LB, Butler A, Martin S, Davies H, Glodzik D, Zou X, Ramakrishna M, Staaf J, Ringnér M, Sieuwerts A, Ferrari A, Morganella S, Fleischer T, Kristensen V, Gut M, van de Vijver MJ, Børresen-Dale AL, Richardson AL, Thomas G, Gut IG, Martens JWM, Foekens JA, Stratton MR, Stunnenberg HG. Partially methylated domains are hypervariable in breast cancer and fuel widespread CpG island hypermethylation. Nat. Commun. 10, 1749 (2019).

19. Fox-Fisher I, Piyanzin S, Ochana BL, Klochendler A, Magenheim J, Peretz A, Loyfer N, Moss J, Cohen D, Drori Y, Friedman N, Mandelboim M, Rothenberg ME, Caldwell JM, Rochman M, Jamshidi A, Cann G, Lavi D, Kaplan T, Glaser B, Shemer R, Dor Y. Remote immune processes revealed by immune-derived circulating cell-free DNA. eLife 10, (2021).

20. Tsui DWY, Cheng ML, Shady M, Yang JL, Stephens D, Won H, Srinivasan P, Huberman K, Meng F, Jing X, Patel J, Hasan M, Johnson I, Gedvilaite E, Houck-Loomis B, Socci ND, Selcuklu SD, Seshan VE, Zhang H, Chakravarty D, Zehir A, Benayed R, Arcila M, Ladanyi M, Funt SA, Feldman DR, Li BT, Razavi P, Rosenberg J, Bajorin D, Iyer G, Abida W, Scher HI, Rathkopf D, Viale A, Berger MF, Solit DB. Tumor fraction-guided cell-free DNA profiling in metastatic solid tumor patients. Genome Med. 13, 96 (2021).

21. Husain H, Pavlick DC, Fendler BJ, Madison RW, Decker B, Gjoerup O, Parachoniak CA, McLaughlin-Drubin M, Erlich RL, Schrock AB, Frampton GM, Das Thakur M, Oxnard GR, Tukachinsky H. Tumor Fraction Correlates With Detection of Actionable Variants Across > 23,000 Circulating Tumor DNA Samples. JCO Precis. Oncol. 6, e2200261 (2022).

22. Beltran H, Romanel A, Conteduca V, Casiraghi N, Sigouros M, Franceschini GM, Orlando F, Fedrizzi T, Ku SY, Dann E, Alonso A, Mosquera JM, Sboner A, Xiang J, Elemento O, Nanus DM, Tagawa ST, Benelli M, Demichelis F. Circulating tumor DNA profile recognizes transformation to castration-resistant neuroendocrine prostate cancer. J. Clin. Invest. 130, 1653–1668 (2020).

23. Prandi D, Baca SC, Romanel A, Barbieri CE, Mosquera JM, Fontugne J, Beltran H, Sboner A, Garraway LA, Rubin MA, Demichelis F. Unraveling the clonal hierarchy of somatic genomic aberrations. Genome Biol. 15, 439 (2014).

24. Lindström LS, Karlsson E, Wilking UM, Johansson U, Hartman J, Lidbrink EK, Hatschek T, Skoog L, Bergh J. Clinically used breast cancer markers such as estrogen receptor, progesterone receptor, and human epidermal growth factor receptor 2 are unstable throughout tumor progression. J. Clin. Oncol. Off. J. Am. Soc. Clin. Oncol. 30, 2601–8 (2012).

25. Morganti, S., Parsons, H. A., Lin, N. U. & Grinshpun, A. Liquid biopsy for brain metastases and leptomeningeal disease in patients with breast cancer. NPJ Breast Cancer 9, 43 (2023).

26. Teschendorff, A. E., Breeze, C. E., Zheng, S. C. & Beck, S. A comparison of reference-based algorithms for correcting cell-type heterogeneity in Epigenome-Wide Association Studies. BMC Bioinformatics 18, 105 (2017).

27. De Luca F, Rotunno G, Salvianti F, Galardi F, Pestrin M, Gabellini S, Simi L, Mancini I, Vannucchi AM, Pazzagli M, Di Leo A, Pinzani P. Mutational analysis of single circulating tumor cells by next generation sequencing in metastatic breast cancer. Oncotarget 7, 26107–26119 (2016).

28. Galardi F, De Luca F, Biagioni C, Migliaccio I, Curigliano G, Minisini AM, Bonechi M, Moretti E, Risi E, McCartney A, Benelli M, Romagnoli D, Cappadona S, Gabellini S, Guarducci C, Conti V, Biganzoli L, Di Leo A, Malorni L. Circulating tumor cells and palbociclib treatment in patients with ER-positive, HER2-negative advanced breast cancer: results from a translational sub-study of the TREnd trial. Breast Cancer Res. BCR 23, 38 (2021).

29. Pestrin M, Salvianti F, Galardi F, De Luca F, Turner N, Malorni L, Pazzagli M, Di Leo A, Pinzani P. Heterogeneity of PIK3CA mutational status at the single cell level in circulating tumor cells from metastatic breast cancer patients. Mol. Oncol. 9, 749–757 (2015).

30. Hoadley KA, Yau C, Hinoue T, Wolf DM, Lazar AJ, Drill E, Shen R, Taylor AM, Cherniack AD, Thorsson V, Akbani R, Bowlby R, Wong CK, Wiznerowicz M, Sanchez-Vega F, Robertson AG, Schneider BG, Lawrence MS, Noushmehr H, Malta TM; Cancer Genome Atlas Network; Stuart JM, Benz CC, Laird PW. Cell-of-Origin Patterns Dominate the Molecular Classification of 10,000 Tumors from 33 Types of Cancer. Cell 173, 291–304.e6 (2018).

31. Li J, Wei L, Zhang X, Zhang W, Wang H, Zhong B, Xie Z, Lv H, Wang X. DISMIR: Deep learning-based noninvasive cancer detection by integrating DNA sequence and methylation information of individual cell-free DNA reads. Brief. Bioinform. 22, bbab250 (2021).

32. Bie F, Wang Z, Li Y, Guo W, Hong Y, Han T, Lv F, Yang S, Li S, Li X, Nie P, Xu S, Zang R, Zhang M, Song P, Feng F, Duan J, Bai G, Li Y, Huai Q, Zhou B, Huang YS, Chen W, Tan F, Gao S. Multimodal analysis of cell-free DNA whole-methylome sequencing for cancer detection and localization. Nat. Commun. 14, 6042 (2023).

33. Schrijver WAME, Suijkerbuijk KPM, van Gils CH, van der Wall E, Moelans CB, van Diest PJ. Receptor Conversion in Distant Breast Cancer Metastases: A Systematic Review and Meta-analysis. J. Natl. Cancer Inst. 110, 568–580 (2018).

34. Shiino S, Ball G, Syed BM, Kurozumi S, Green AR, Tsuda H, Takayama S, Suto A, Rakha EA. Prognostic significance of receptor expression discordance between primary and recurrent breast cancers: a meta-analysis. Breast Cancer Res. Treat. 191, 1–14 (2022).

35. Aftimos P, Oliveira M, Irrthum A, Fumagalli D, Sotiriou C, Gal-Yam EN, Robson ME, Ndozeng J, Di Leo A, Ciruelos EM, de Azambuja E, Viale G, Scheepers ED, Curigliano G, Bliss JM, Reis-Filho JS, Colleoni M, Balic M, Cardoso F, Albanell J, Duhem C, Marreaud S, Romagnoli D, Rojas B, Gombos A, Wildiers H, Guerrero-Zotano A, Hall P, Bonetti A, Larsson KF, Degiorgis M, Khodaverdi S, Greil R, Sverrisdóttir Á, Paoli M, Seyll E, Loibl S, Linderholm B, Zoppoli G, Davidson NE, Johannsson OT, Bedard PL, Loi S, Knox S, Cameron DA, Harbeck N, Montoya ML, Brandão M, Vingiani A, Caballero C, Hilbers FS, Yates LR, Benelli M, Venet D, Piccart MJ. Genomic and Transcriptomic Analyses of Breast Cancer Primaries and Matched Metastases in AURORA, the Breast International Group (BIG) Molecular Screening Initiative. Cancer Discov. 11, 2796–2811 (2021).

36. Zardavas D, Maetens M, Irrthum A, Goulioti T, Engelen K, Fumagalli D, Salgado R, Aftimos P, Saini KS, Sotiriou C, Campbell P, Dinh P, von Minckwitz G, Gelber RD, Dowsett M, Di Leo A, Cameron D, Baselga J, Gnant M, Goldhirsch A, Norton L, Piccart M. The AURORA initiative for metastatic breast cancer. Br. J. Cancer 111, 1881–7 (2014).

37. Hulsbergen AFC, Claes A, Kavouridis VK, Ansaripour A, Nogarede C, Hughes ME, Smith TR, Brastianos PK, Verhoeff JJC, Lin NU, Broekman MLD. Subtype switching in breast cancer brain metastases: a multicenter analysis. Neuro-Oncol. 22, 1173–1181 (2020).

38. Prat A, Brasó-Maristany F, Martínez-Sáez O, Sanfeliu E, Xia Y, Bellet M, Galván P, Martínez D, Pascual T, Marín-Aguilera M, Rodríguez A, Chic N, Adamo B, Paré L, Vidal M, Margelí M, Ballana E, Gómez-Rey M, Oliveira M, Felip E, Matito J, Sánchez-Bayona R, Suñol A, Saura C, Ciruelos E, Tolosa P, Muñoz M, González-Farré B, Villagrasa P, Parker JS, Perou CM, Vivancos A. Circulating tumor DNA reveals complex biological features with clinical relevance in metastatic breast cancer. Nat. Commun. 14, 1157 (2023).

39. Benelli M, Franceschini GM, Magi A, Romagnoli D, Biagioni C, Migliaccio I, Malorni L, Demichelis F. Charting differentially methylated regions in cancer with Rocker-meth. Commun. Biol. 4, 1249 (2021).

40. Zviran A, Schulman RC, Shah M, Hill STK, Deochand S, Khamnei CC, Maloney D, Patel K, Liao W, Widman AJ, Wong P, Callahan MK, Ha G, Reed S, Rotem D, Frederick D, Sharova T, Miao B, Kim T, Gydush G, Rhoades J, Huang KY, Omans ND, Bolan PO, Lipsky AH, Ang C, Malbari M, Spinelli CF, Kazancioglu S, Runnels AM, Fennessey S, Stolte C, Gaiti F, Inghirami GG, Adalsteinsson V, Houck-Loomis B, Ishii J, Wolchok JD, Boland G, Robine N, Altorki NK, Landau DA. Genome-wide cell-free DNA mutational integration enables ultra-sensitive cancer monitoring. Nat. Med. 26, 1114–1124 (2020).

41. Jamshidi A, Liu MC, Klein EA, Venn O, Hubbell E, Beausang JF, Gross S, Melton C, Fields AP, Liu Q, Zhang N, Fung ET, Kurtzman KN, Amini H, Betts C, Civello D, Freese P, Calef R, Davydov K, Fayzullina S, Hou C, Jiang R, Jung B, Tang S, Demas V, Newman J, Sakarya O, Scott E, Shenoy A, Shojaee S, Steffen KK, Nicula V, Chien TC, Bagaria S, Hunkapiller N, Desai M, Dong Z, Richards DA, Yeatman TJ, Cohn AL, Thiel DD, Berry DA, Tummala MK, McIntyre K, Sekeres MA, Bryce A, Aravanis AM, Seiden MV, Swanton C. Evaluation of cell-free DNA approaches for multi-cancer early detection. Cancer Cell 40, 1537–1549.e12 (2022).

42. Liu MC, Oxnard GR, Klein EA, Swanton C, Seiden MV; CCGA Consortium. Sensitive and specific multi-cancer detection and localization using methylation signatures in cell-free DNA. Ann. Oncol. Off. J. Eur. Soc. Med. Oncol. 31, 745–759 (2020).

43. Danecek P, Bonfield JK, Liddle J, Marshall J, Ohan V, Pollard MO, Whitwham A, Keane T, McCarthy SA, Davies RM, Li H. Twelve years of SAMtools and BCFtools. GigaScience 10, giab008 (2021).

44. Ewels PA, Peltzer A, Fillinger S, Patel H, Alneberg J, Wilm A, Garcia MU, Di Tommaso P, Nahnsen S. The nf-core framework for community-curated bioinformatics pipelines. Nat. Biotechnol. 38, 276–278 (2020).

45. Krueger, F. & Andrews, S. R. Bismark: A flexible aligner and methylation caller for Bisulfite-Seq applications. Bioinformatics 27, 1571–2 (2011).

