## Supplementary Text and Figures to METER manuscript for "Sensitive tumor detection, accurate quantification, and cancer subtype classification using low-pass whole methylome sequencing of plasma DNA"

#### Preliminary study to ensure the applicability of ichorCNA in low-pass whole genome bisulfite sequencing (lpWGBS) data

We conducted a preliminary analysis to investigate whether pretreatment with bisulfite prior to library construction for low-pass whole genome sequencing (lpWGS) could affect the estimation of DNA copy number alterations (CNA). To this end, we considered 9 Whole Genome Bisulfite Sequencing (WGBS) samples from plasma of patients with advanced prostate cancer and 9 controls, obtained from Beltran et al., 2020[1]. Having an average coverage of 30X, the Whole Genome Bisulfite Sequencing (WGBS) data was initially downsampled to simulate 0.1X coverage (low-pass WGBS, lpWGBS). Subsequently, ichorCNA tool[2] was employed on these subsampled data, using the information from the 9 available control samples as a reference "panel of normals". We then compared the CNA profiles estimated by ichorCNA from lpWGBS to the reference CNA profiles estimated by FACETS[3] from high-coverage (200X) Whole-Exome Sequencing (WES) of the same plasma samples. As shown in **Supplementary Figure 1**, we observed excellent concordance between the CNA profiles estimated from high-coverage WES and those estimated from lpWGBS. Using tumor burden estimates generated by CLONET[4] from WES data as reference, we observed a good consistency between the tumor content estmates derived from lpWGBS by ichorCNA and the expected values (R=0.7, RMSE=0.2) (**Supplementary Figure 2A**). Finally, we analyzed the estimated tumor burden by ichorCNA in controls and found it to be nearly zero (**Supplementary Figure 2B**). These analyses collectively suggest that bisulfite treatment prior to library construction for WGS does not introduce significant biases in CNA estimation, and therefore does not hinder the applicability of ichorCNA on this type of data.

**Microarray-based analysis**

To test the reliability of METER exploiting specific Differentially Methylated Sites (DMS) and Regions (DMR) retrieved from microarray data, we considered the Illumina Infiunium450K array data from the TCGA-BRCA dataset[5]. Rocker-meth[6] was applied to the beta values from TCGA-BRCA dataset, comprising 782 breast tumors, and to the beta values of 380 healthy whole blood samples downloaded from EWAS Data Hub (https://doi.org/10.1093/nar/gkz840). Subsequently, similarly to what done for WGBS-based iDMS and iDMR, we applied a series of filtering criteria to select tumor content (TC) informative DMS and DMR (iDMS and iDMR).

For iDMS, the following criteria were used:

- Area under curve (AUC) of a receiver operating characteristic (ROC) curve, separating tumor from control samples, higher than 0.80 (hyper-methylated sites) or lower than 0.20 (hypo-methylated sites);
- Difference of mean beta values between tumor and control samples greater than 0.40 (where beta value is defined as the per-site proportion of methylated signal over the total methylation signal);
- Third quartile of beta values in control samples lower than 0.1 (hyper-methylated) or first quartile of beta values in control samples above 0.9 (hypo-methylated);

leading to a selection of 3029 hyper and 3302 hypo iDMS.

For iDMR, the following criteria were used:

- FDR<0.05;
- Difference between mean of beta by DMR values between tumor and control samples greater than 20 (where beta by DMR values are computed as the mean beta values of CpG sites within the specific DMR);

leading to a selection of 705 hyper and 738 hypo iDMR.

Importantly, in the selection of array-based DMR, we omitted filters based on the beta distribution of control healthy blood samples, maintaining the essential filters reported above. This decision was made to account for the limited number of DMR obtained from array data compared to those from WGBS data.

**TC quantification using METER-quant based on array iDMS**

As for METER-quant with WGBS-based iDMS, the proportion of tumor-like sites (PTS) for each sample was computed (that is the ratio of hypermethylated iDMS with beta=100% or hypomethyalted iDMS with beta=0% over the total iDMS with beta=100% covered) as a proxy of sample's TC. Only reads with alpha value of 100% were considered for PTS calculation. As shown in **Supplementary Figure 9A,** we observed TC values in mBC cfDNA samples (N=124) significantly higher than TC values in control cfDNA samples (N=30) within the study cohort. Furthermore, we observed high concordance between TC estimates by METER-quant with array-based iDMS and TC estimates by ichorCNA on the mBC cfDNA samples within the study cohort (**Supplementary Figure 9B**).

**TC detection using METER detect based on array iDMR**

As for METER-detect with WGBS-based iDMR, the proportion of tumor-like reads (PTR) for each sample was computed considering only reads with alpha value of 100% (that is the proportion of fully methylated or unmethylated reads within selected hyper- or hypo- iDMR respectively over the total reads with alpha value of 100%). Regarding the minimum number of CpG sites comprised in the reads, in this case we required a minimum of 4 CpG sites (accounting for the reduced number of DMR retrieved, compared to WGBS-based DMR). Following this, a Z-score method was employed, leveraging the distribution of this measure within control samples (reference model), to categorize each sample. If a sample's Z-score exceeded a threshold derived from the Z-scores computed in control samples, it was classified as ctDNA positive (METER+); otherwise, it was classified as ctDNA negative (METER-). Considering only samples with TC by ichorCNA<0.03 (i.e. undetected by ichorCNA, ichorCNA-) within our study cohort (N=60 tumors and N=30 controls), PTR of tumor cfDNA samples were significantly higher of PTR of control samples, and using a threshold corresponding to Z-score=3, 43% of ichorCNA- samples were classified as METER+ (**Supplementary Figure 9D**). We observed globally higher distributions of PTR both in control and in tumor samples, compared to those obtained by METER-detect with WGBS-based iDMR. This result is expected, considering the lack of constraints imposed on beta distribution of healthy plasma samples employed for iDMR selection from array data. Similarly to what done for METER-detect with WGBS-based DMR, using a leave-one-out strategy applied to control samples, we evaluated FDR at different Z-score thresholds, obtaining FDR<5% for Z-score=3 (**Supplementary Figure 9E**).

**Association of patients' outcome with METER-detect classification obtained using**

**Array-based DMR detected by Rocker-meth in TCGA-BRCA dataset**

We evaluated the efficacy of METER-detect with array-based iDMR in stratifying patients in our study cohort based on their association with Progression Free Survival (PFS) and Overall Survival (OS). Regarding OS, METER+ patients exhibited significantly shorter survival times compared to METER- counterparts. However, among ichorCNA- patients (TC by ichorCNA<0.03, n=23), no substantial difference was noted (**Supplementary Figure 11A**). Regarding PFS, METER+ patients showed poorer outcomes compared to METER-, while no significant association was found with ichorCNA classification. Among ichorCNA- patients (n=23), although not statistically significant, we observed a trend, although not significant, for METER+ patients to have worse PFS (**Supplementary Figure 11B**).
To evaluate the dynamic changes in ctDNA from baseline (BL) to Cycle 2 Day 1 (C2D1), we stratified patients based on METER-detect and ichorCNA at C2D1 versus BL (METER or ichorCNA +/+, +/-, -/-), excluding patients where ctDNA was detected only at C2D1 due to limited observations. METER-detect demonstrated comparable ability to ichorCNA in stratifying patients for both OS and PFS. Among ichorCNA -/- patients (n=15), no significant stratification was observed for OS, while, for PFS, METER-detect showed a trend towards significant stratification (**Supplementary Figure 11C and 11D**).

**Considerations on the use of array-based iDMR and iDMS**

Using array-based DMS and DMR we observed consistent findings for METER-quant and METER-detect with those observed using WGBS-based DMS and DMR. However, we observed poorer performance for METER-detect with array-based DMR, particularly in its ability to stratify patients based on PFS and OS. These findings, in particular those regarding METER-detect, are expected due to the lower number of array-based DMR compared to those identified from WGBS. Given these low coverages and therefore limited number of reads per sample, a greater number of DMR coupled with more stringent filters in DMR selection would be necessary to achieve higher sensitivity and specificity. To address this issue with array-based DMR, it may be beneficial to increase coverage of lpWGBS data to analyze, to increase the number of reads per sample and thereby increase the signal-to-noise ratio. However, further analyses would be essential to validate this proposed approach.

### References


1. Beltran H, Romanel A, Conteduca V, Casiraghi N, Sigouros M, Franceschini GM, Orlando F, Fedrizzi T, Ku SY, Dann E, Alonso A, Mosquera JM, Sboner A, Xiang J, Elemento O, Nanus DM, Tagawa ST, Benelli M, Demichelis F. Circulating tumor DNA profile recognizes transformation to castration-resistant neuroendocrine prostate cancer. J. Clin. Invest. 130, 1653–1668 (2020).

2. Adalsteinsson VA, Ha G, Freeman SS, Choudhury AD, Stover DG, Parsons HA, Gydush G, Reed SC, Rotem D, Rhoades J, Loginov D, Livitz D, Rosebrock D, Leshchiner I, Kim J, Stewart C, Rosenberg M, Francis JM, Zhang CZ, Cohen O, Oh C, Ding H, Polak P, Lloyd M, Mahmud S, Helvie K, Merrill MS, Santiago RA, O'Connor EP, Jeong SH, Leeson R, Barry RM, Kramkowski JF, Zhang Z, Polacek L, Lohr JG, Schleicher M, Lipscomb E, Saltzman A, Oliver NM, Marini L, Waks AG, Harshman LC, Tolaney SM, Van Allen EM, Winer EP, Lin NU, Nakabayashi M, Taplin ME, Johannessen CM, Garraway LA, Golub TR, Boehm JS, Wagle N, Getz G, Love JC, Meyerson M. Scalable whole-exome sequencing of cell-free DNA reveals high concordance with metastatic tumors. Nat. Commun. 8, 1324 (2017).

3. Shen R, Seshan VE. FACETS: allele-specific copy number and clonal heterogeneity analysis tool for high-throughput DNA sequencing. Nucleic Acids Res. (2016).

4. Prandi D, Baca SC, Romanel A, Barbieri CE, Mosquera JM, Fontugne J, Beltran H, Sboner A, Garraway LA, Rubin MA, Demichelis F. Unraveling the clonal hierarchy of somatic genomic aberrations. Genome Biol. 15, 439 (2014).

5. Hoadley KA, Yau C, Hinoue T, Wolf DM, Lazar AJ, Drill E, Shen R, Taylor AM, Cherniack AD, Thorsson V, Akbani R, Bowlby R, Wong CK, Wiznerowicz M, Sanchez-Vega F, Robertson AG, Schneider BG, Lawrence MS, Noushmehr H, Malta TM; Cancer Genome Atlas Network; Stuart JM, Benz CC, Laird PW. Cell-of-Origin Patterns Dominate the Molecular Classification of 10,000 Tumors from 33 Types of Cancer. Cell 173, 291-304.e6 (2018).

6. Benelli M, Franceschini GM, Magi A, Romagnoli D, Biagioni C, Migliaccio I, Malorni L, Demichelis F. Charting differentially methylated regions in cancer with Rocker-meth. Commun. Biol. 4, 1249 (2021).

7. Doebley AL, Ko M, Liao H, Cruikshank AE, Santos K, Kikawa C, Hiatt JB, Patton RD, De Sarkar N, Collier KA, Hoge ACH, Chen K, Zimmer A, Weber ZT, Adil M, Reichel

JB, Polak P, Adalsteinsson VA, Nelson PS, MacPherson D, Parsons HA, Stover DG, Ha G. A framework for clinical cancer subtyping from nucleosome profiling of cell-free DNA. Nat. Commun. 13, 7475 (2022).

### Supplementary Figures

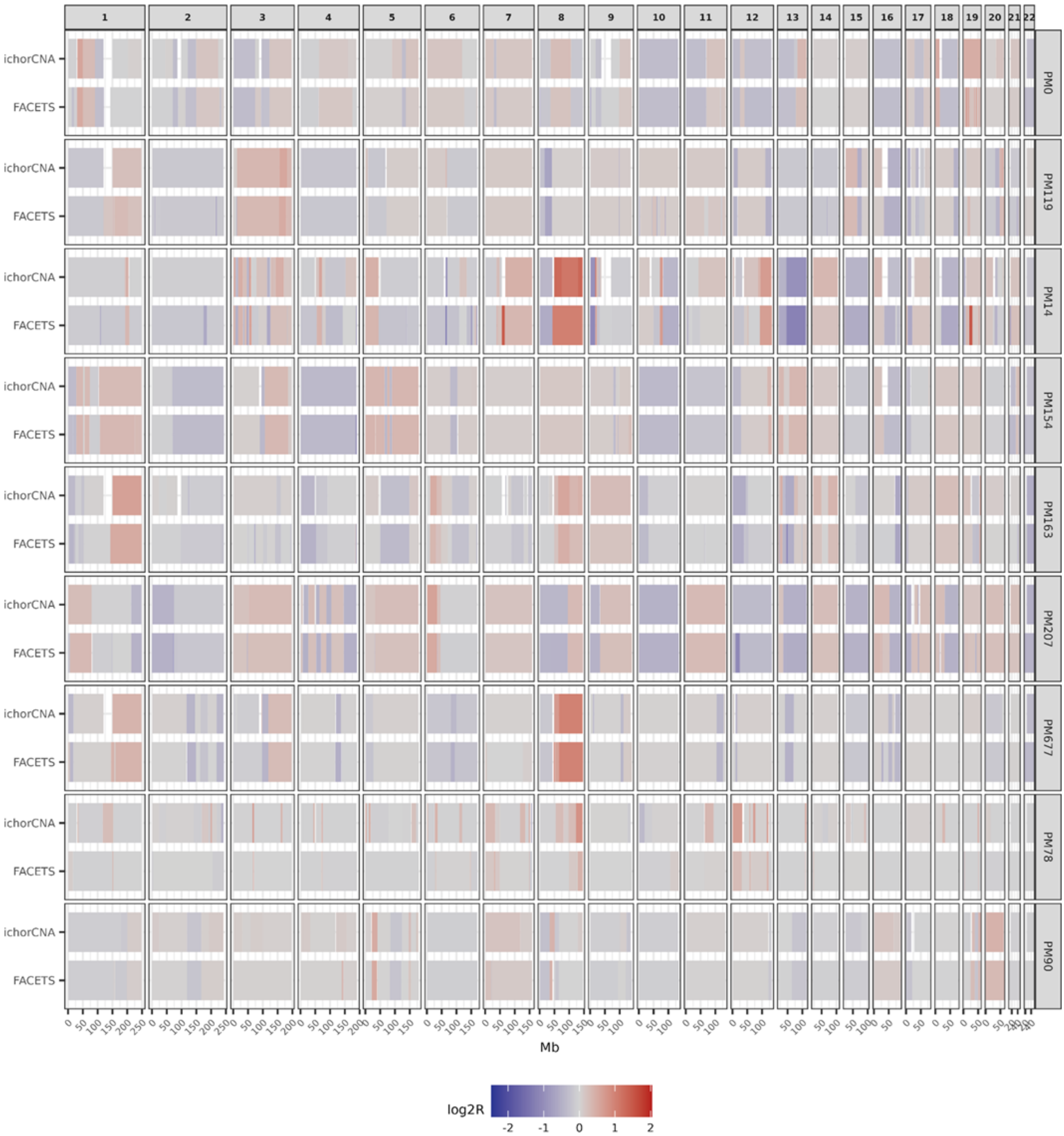


**Fig. S1: Preliminary study to ensure the applicability of ichorCNA in lpWGBS data: CNA profiles.** Segment plots showing the concordance of mean log2ratio values along the genome (genomic coordinates on the x-axis) estimated by ichorCNA from lpWGBS data and by FACETS from high coverage WES data of 9 cfDNA samples of advanced prostate cancer patients from Beltran et al., 2020. IchorCNA was applied using available controls as reference PoN and 1Mb window size. log2ratio: logarithm base 2 of estimated tumor signal compared to expected signal in a copy number neutral genome; Mb: mega bases; lpWGBS: low-pass whole genome bisulfite sequencing data; WES: whole exome sequencing; PoN: panel of normals.

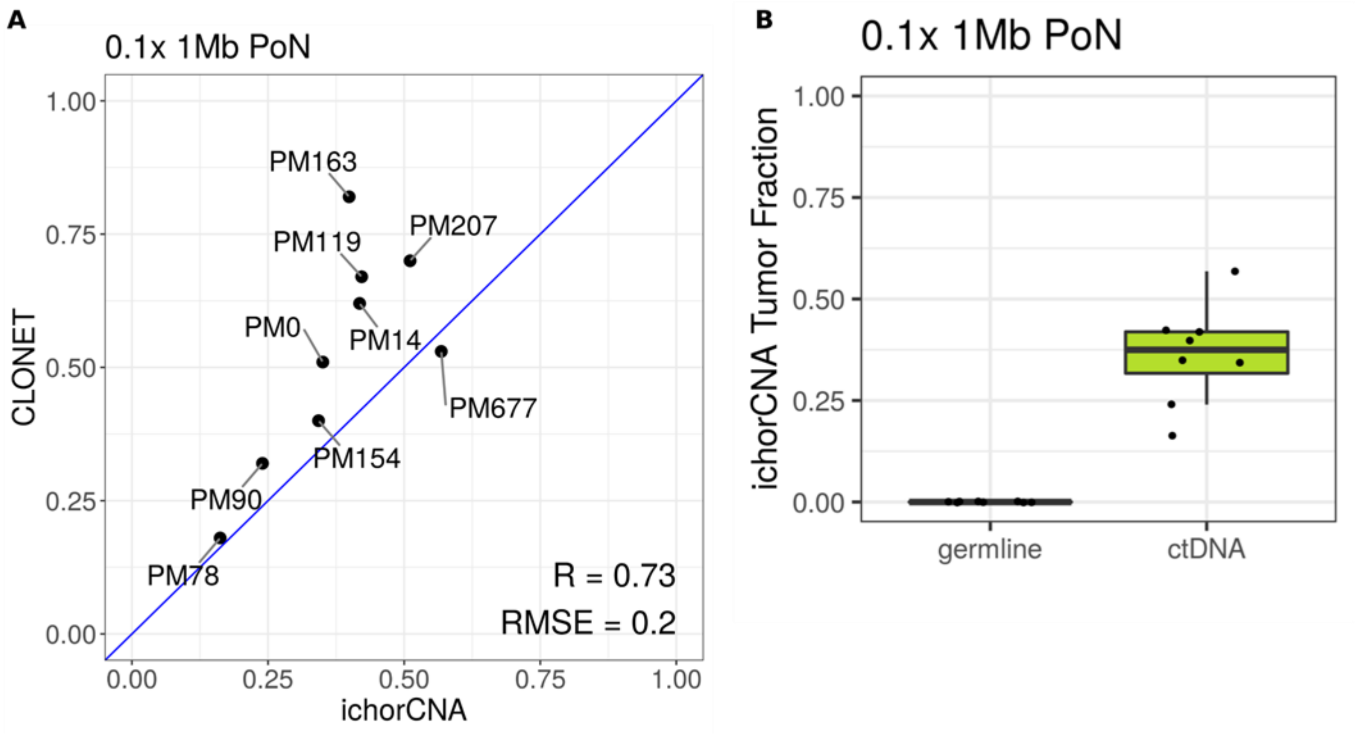


**Fig. S2: Preliminary study to ensure the applicability of ichorCNA in lpWGBS data: TC estimates.** (A) Scatter plot showing the concordance between tumor fraction estimates by CLONET from WES (y-axis) and by ichorCNA from lpWGBS data (x-axis) of 9 cfDNA samples of advanced prostate cancer patients from Beltran et al., 2020. (B) Box plots showing the distribution of TC estimates by ichorCNA in advanced prostate cancer and matched control plasma samples (N=9) from Beltran et al., 2020. IchorCNA was applied using available controls as reference PoN and 1Mb window size. Mb: mega bases; PoN: panel of normals; R: Pearson's correlation coefficient; RMSE: root mean squared error; germline: control samples; ctDNA: circulating tumor DNA corresponding to advanced prostate cancer plasma samples; lpWGBS: low-pass whole genome bisulfite sequencing data; WES: whole exome sequencing; cfDNA: cell-free DNA; TC: tumor content.

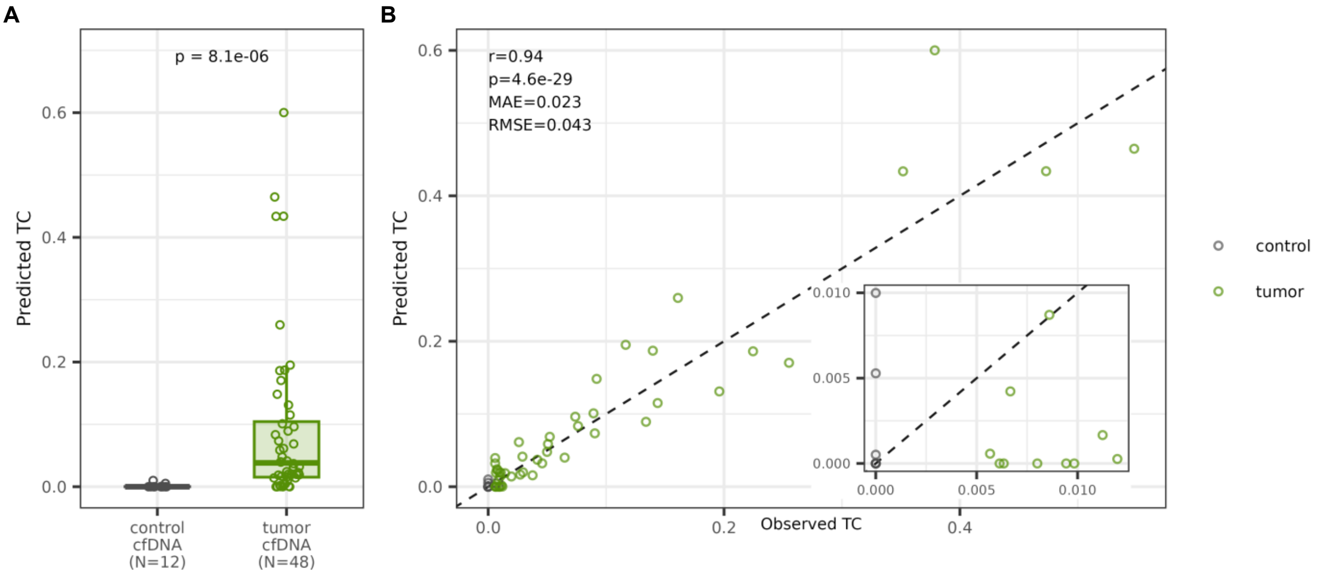


**Fig. S3: Linear model on test set.** (A) Box plot showing the distribution of TC estimates by the linear model in control and mBC cfDNA samples within the test set (N=60 (40%) samples, N=48 tumors and N=12 controls). P-value was estimated by Wilcoxon test (B) Scatter plot showing the concordance between observed TC (ichorCNA estimates for tumor samples and 0 for controls) and predicted TC on the test set. The inset plot reports samples exhibiting predicted TC within the range of predicted TC in controls. R and p are Pearson's correlation coefficient and corresponding p-value, respectively. cfDNA: cell-free DNA; TC: tumor content, MAE: mean absolute error, RMSE, root mean squared error.

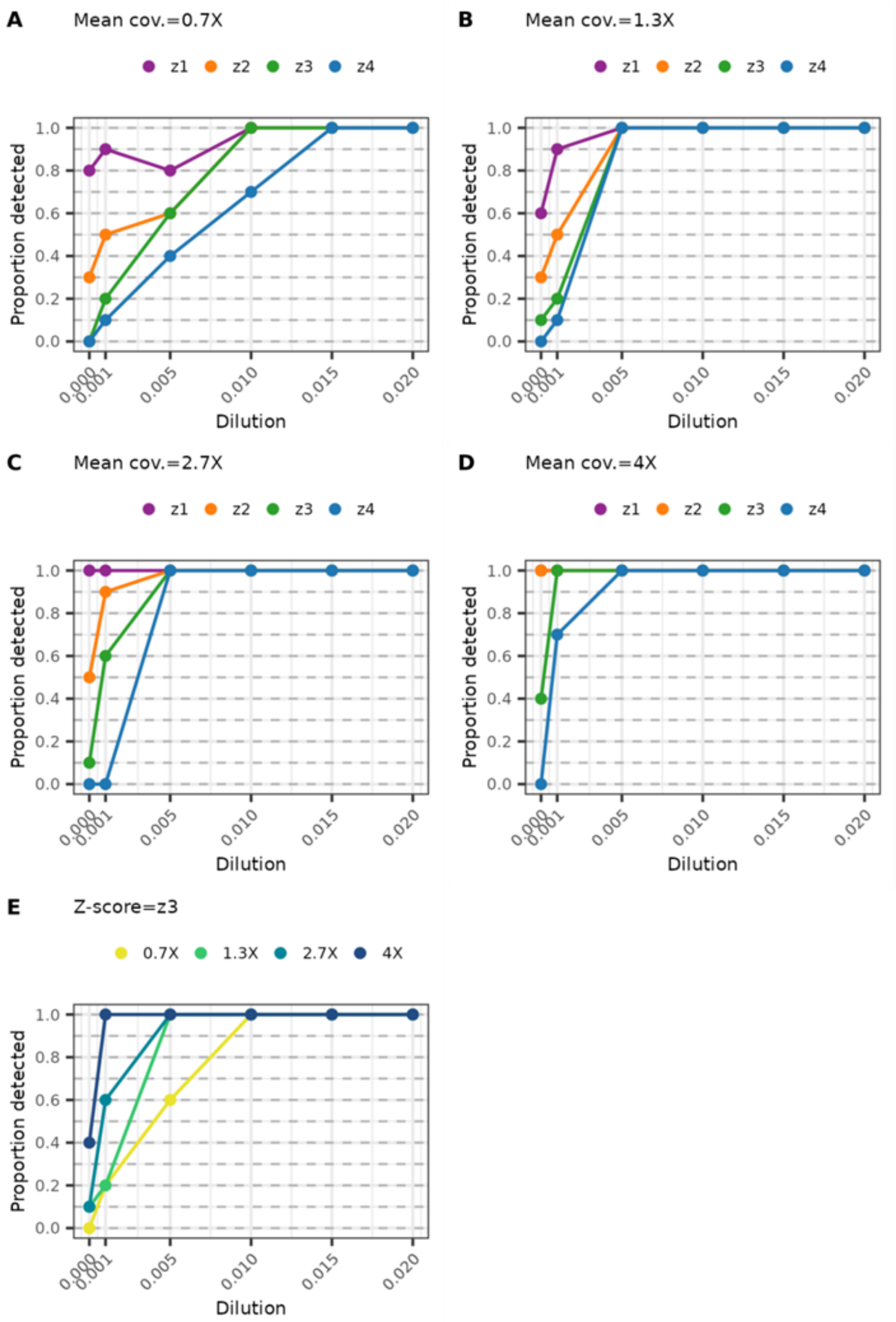


**Fig. 4: in-silico studies.** (A), (B), (C) and (D) Line plots showing in-silico dilutions study to evaluate detection sensitivity and FDR of METER-detect at increasing mean coverage levels ((A) to (D)) and Z-score thresholds. (E) Line plot showing detection sensitivity and FDR of METER-detect at increasing coverage using a threshold corresponding to Z-score=3. FDR: false discovery rate; TC: tumor content.

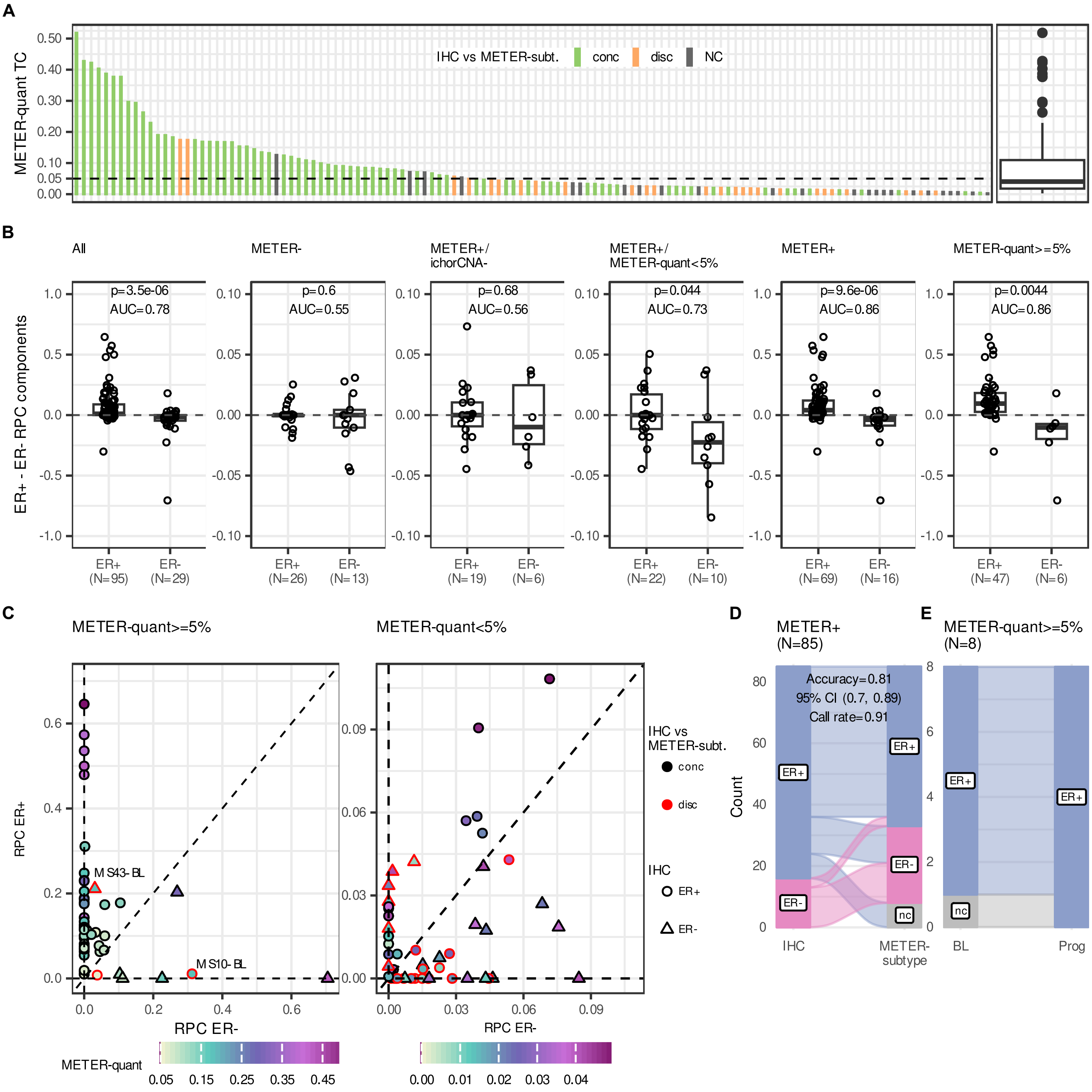


**Fig. S5: ER subtyping from cfDNA samples by METER-subtype.** (A) Bar plot and box plot showing the distribution of samples' TC by METER quant (y-axis) and concordance between ER subtyping by IHC and METER-subtype. The dashed line indicates METER-quant=5%. (B) Box plots showing the distribution of the difference between ER+ and ER- RPC components in ER+ and ER- BC samples, as classified by IHC, within the study cohort. P is Wilcoxon Test p-value. (C) Scatter plots showing ER+ (y-axis) and ER- (x-axis) RPC components for each sample with TC by METER-quant>=5% (left) and with TC by METER-quant<5% (right). ER status according to IHC, concordance between ER subtyping by IHC and METER-subtype, and TC by

METER-quant are shown for each sample. (D) Alluvial plot showing concordance between ER subtyping by IHC and METER-subtype on METER+ cfDNA samples within the study cohort. (E) Alluvial plot showing concordance between ER subtyping by METER-subtype at BL and Prog for the 8 patients with available BL/Prog samples and METER-quant>=5% in both time points (all ER+ based on IHC). IHC: immunohistochemistry; conc: concordant ER status by IHC and METER-subtype; disc: discordant ER status by IHC and METER-subtype; NC: not classified by METER-subtype; RPC: robust partial correlation; AUC: area under the curve; CI: confidence interval; BL: baseline; Prog: progression of disease; TC: tumor content; cfDNA: cell-free DNA.

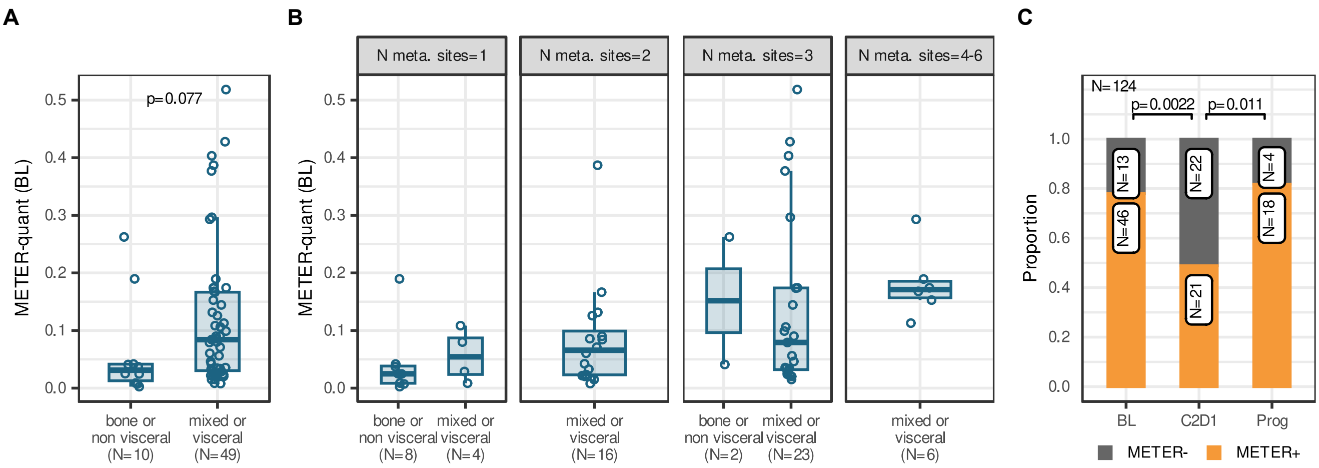


**Fig. 6: Association of METER results with clinically relevant prognostic factors.** (A) and (B) Box plots showing the distribution of TC by METER-quant in mBC samples at BL (N=59) stratified by visceral and bone or non-visceral disease at study entry (A) and by number of metastatic sites combined with visceral and bone or non-visceral disease at study entry (B). P is Wilcoxon Test p-value. (C) Bar plot showing the proportion of METER+ samples (y-axis) across clinical time points (x-axis) for all cfDNA samples at BL (N=59), at C2D1 (N=43) and at Prog (N=22) within the study cohort. P is Two-Proportions z-Test p-value. TC: tumor content; mBC: metastatic Breast Cancer; cfDNA: cell-free DNA. BL: baseline; C2D1: cycle 2 day 1; Prog: progression of disease.

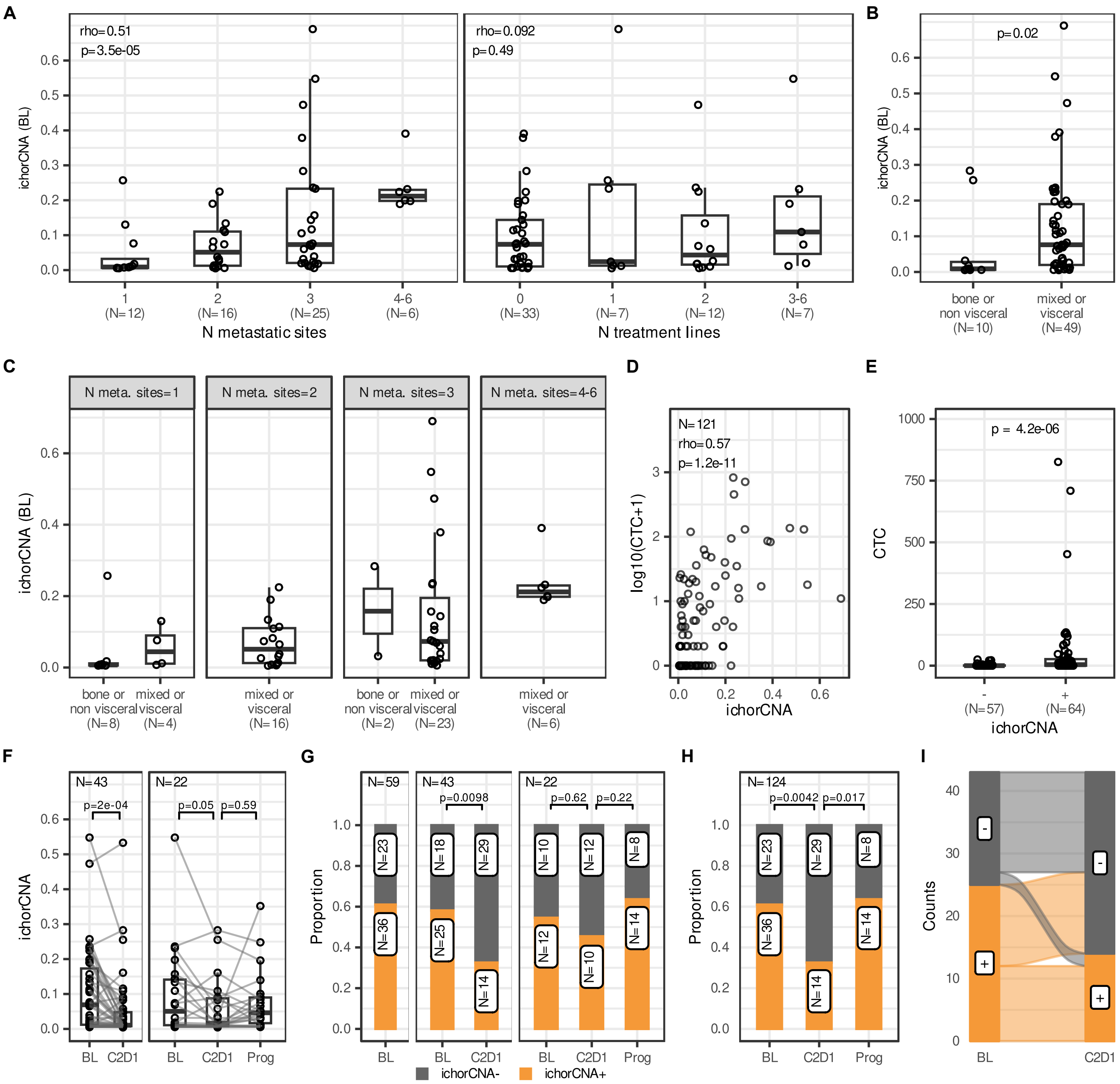


**Fig. S7: Association of ichorCNA results with clinically relevant prognostic factors.** (A) Box plots showing the distribution of TC by ichorCNA in mBC cfDNA samples at BL (N=59) stratified by number of metastatic sites at study entry (left) and number of metastatic treatment lines at study entry (right). p is Pearson's correlation Test p-value. (B) and (C) Box plots showing the distribution of TC by ichorCNA in mBC cfDNA samples at BL (N=59) stratified by visceral and bone or non-visceral disease at study entry (B) and by number of metastatic sites combined with visceral and bone or non-visceral disease at study entry (C). P is Wilcoxon Test p-value. (D) Scatter plot of TC by ichorCNA (x-axis) versus the logarithm base 10 of the number of CTC in samples (y-axis). P is Pearson's correlation Test p-value. (E) Box plots showing the distribution of the number of CTC in ichorCNA- and ichorCNA+ samples, for all cfDNA samples (N=124) within the study cohort. P is Wilcoxon Test p-value. (F) Box plots showing the distribution of samples' TC by ichorCNA (y-axis) across clinical time points (x-axis) for patients having both BL and C2D1 time points (left panel, N=43) and for patients with complete longitudinal data (right panel, N=22) within the study cohort. P is paired Wilcoxon p-value. (G) Bar plots showing the proportion of ichorCNA+ cfDNA samples (y-axis) across clinical time points (x-axis) for all patients at BL (left panel, N=59), for patients having both BL and C2D1 time points (middle panel, N=43) and for patients with complete longitudinal data (right panel, N=22) within the study cohort. P is McNemar's chi-squared Test p-value. (H) Bar plot showing the proportion of ichorCNA+ samples (y-axis) across clinical time points (x-axis) for all cfDNA samples at BL (N=59), at C2D1 (N=43) and at Prog (N=22) within the study cohort. P is Two-Proportions z-Test p-value. (I) Alluvial plot showing the dynamics of samples' classification by ichorCNA in patients having both BL and C2D1 time points (N=43) within the study cohort. Rho: Spearman's correlation coefficient; TC: tumor content; mBC: metastatic Breast Cancer; cfDNA: cell-free DNA; CTC: circulating tumor cells; ichorCNA-: TC<3% according to ichorCNA; ichorCNA+: TC>=3% according to ichorCNA; BL: pre-treatment baseline; C2D1:cycle 2 day 1; Prog: progression.

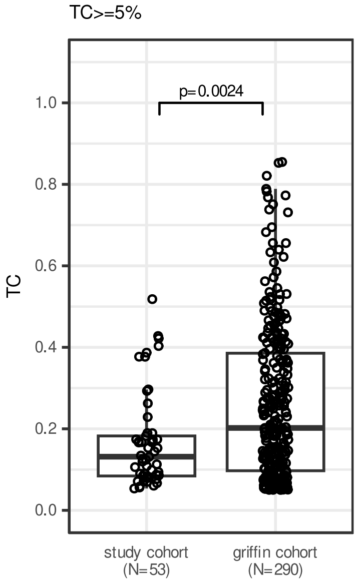


**Fig. S8: TC distribution of mBC cfDNA samples in MIMESIS and the GRIFFIN study cohort (Doebley et al., 2022).** Box plots showing the distribution of TC of mBC cfDNA samples within the MIMESIS study cohort and study cohort from Doebley et al. 2022, for samples with TC>=5%. TC for MIMESIS lpWGBS study cohort has been obtained by METER-quant, TC for the study cohort from Dobley et al. 2022 has been obtained by ichorCNA. P is Wilcoxon Rank Sum Test p-values. TC: tumor content, mBC: metastatic breast cancer.

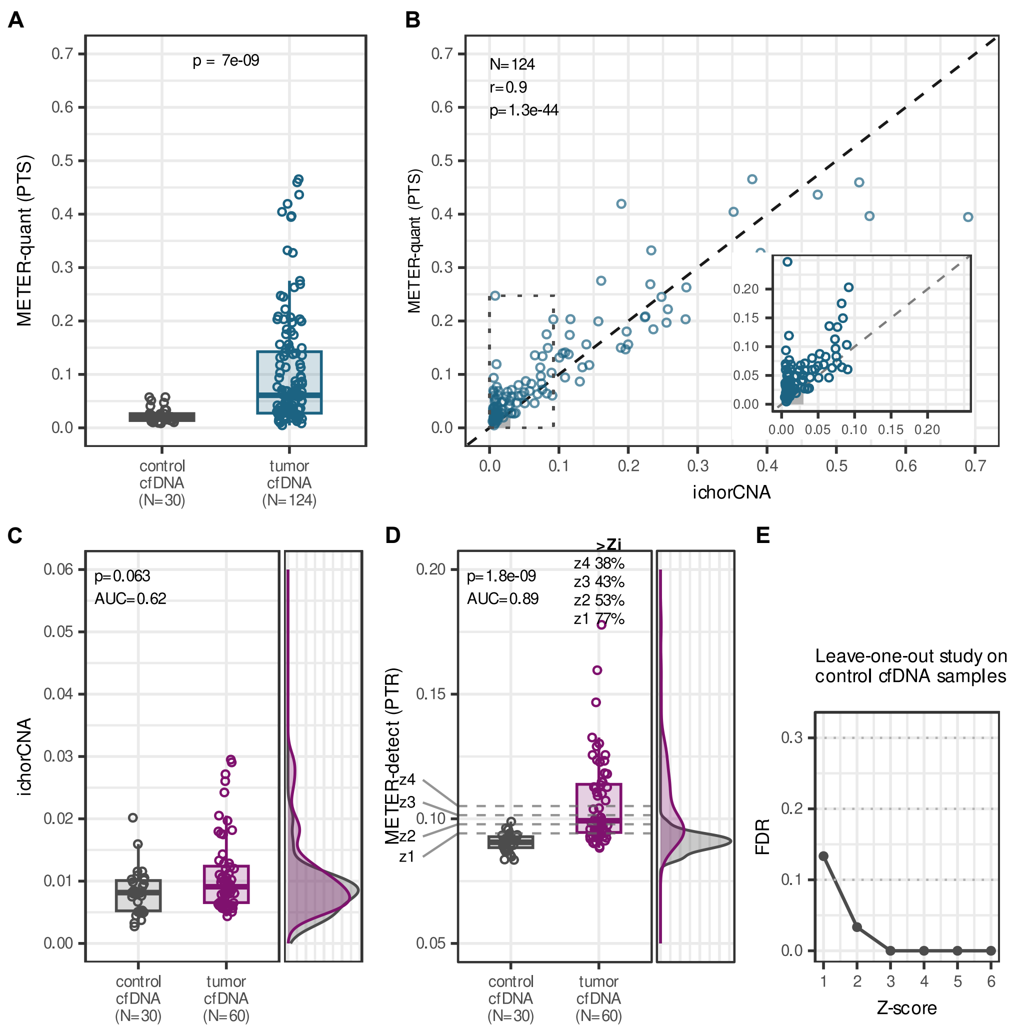


**Fig. S9: TC quantification by METER-quant and TC detection by METER-detect using array-based DMR detected by Rocker-meth in TCGA-BRCA dataset.** (A) Boxplots showing the distribution of TC estimates by METER-quant using array-based iDMS in control and mBC cfDNA samples within the study cohort. P-value was estimated by Wilcoxon test (B) Scatter plot showing TC estimates of mBC cfDNA samples by METER-quant (y-axis) and ichorCNA (x-axis). The inset plot reports samples with TC range of 0-10% based on ichorCNA. The shaded region highlights the range of TC where ichorCNA is not applicable (0-3%). R and p are Pearson's correlation coefficient and corresponding p-value, respectively. (C) and (D) Boxplots and density plots showing the distribution of TC by ichorCNA (C) and PTR by METER-detect with array-based iDMR (D) in 30 control and 60 mBC cfDNA samples within the study cohort with TC by ichorCNA<3%. In (D), the dashed lines indicate different Z-score thresholds based on control samples' PTR distribution and percentages of tumor samples with Z-scores exceeding Z-score thresholds (Zi) are shown. P is p-value estimated by Wilcoxon test. (E) Leave-one-out study on control cfDNA samples to estimate FDR of METER-detect at different Z-score thresholds. PTS, proportion of tumor-like sites; r: Pearson's correlation coefficient; rho: Spearman's correlation coefficient; PTR: proportion of tumor-like reads; AUC: area under the curve; FDR: false discovery rate; TC: tumor content; mBC: metastatic breast cancer; cfDNA: cell-free DNA.

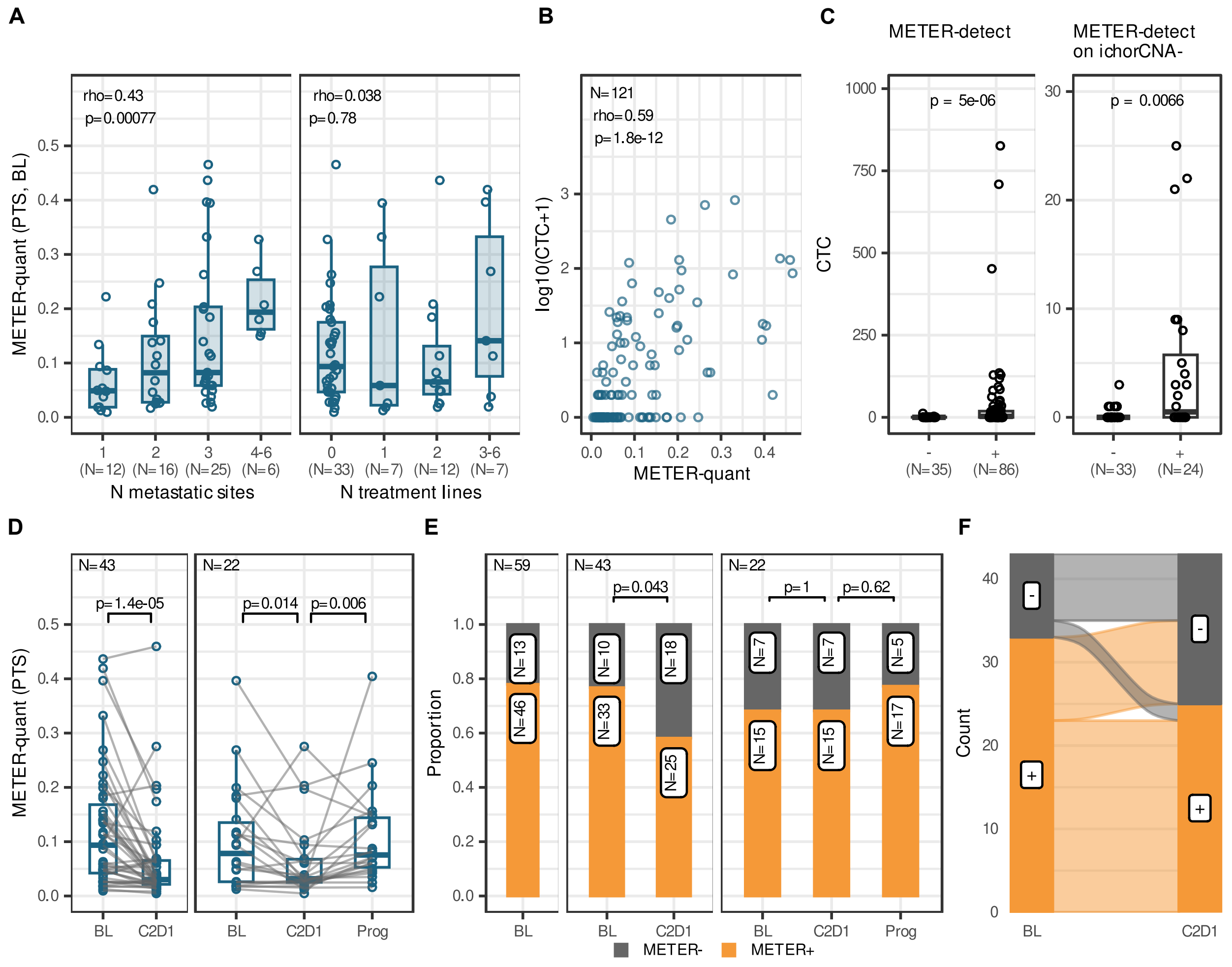


**Fig. S10: Association of clinically relevant prognostic factors with METER.** estimates using array-based DMR detected by Rocker-meth in TCGA-BRCA dataset (A) Box plots showing the distribution of TC by METER-quant in mBC samples at BL (N=59) stratified by number of metastatic sites at study entry (left) and number of metastatic treatment lines at study entry (right). p is Pearson's correlation Test p-value. (B) Scatterplot of TC by METER-quant (x-axis) versus the logarithm base 10 of the number of CTC in samples (y-axis). p is Pearson's correlation Test p-value. (C) Box plots showing the distribution of the number of CTC in METER- and METER+ samples, for all samples (N=124, left panel) and for samples with TC by ichorCNA<3% (N=57, right panel) within the study cohort. p is Wilcoxon Test p-value. (D) Box plots showing the distribution of samples' TC by METER-quant (y-axis) across clinical time points (x-axis) for patients having both BL and C2D1 time points (left panel, N=43) and for patients with complete longitudinal data (right panel, N=22) within the study cohort. p is paired Wilcoxon p-value. (E) Bar plots showing the proportion of METER+ cfDNA samples (y-axis) across clinical time points (x-axis) for all patients at BL (left panel, N=59), for patients having both BL and C2D1 time points (middle panel, N=43) and for patients with complete longitudinal data (right panel, N=22) within the study cohort. p is McNemar's chi-squared Test p-value. (F) Alluvial plot showing the dynamics of samples' classification by METER-detect in patients having both BL and C2D1 time points (N=43) within the study cohort. Rho: Spearman's correlation coefficient; TC: tumor content; CTC: circulating tumor cells; BL: pre-treatment baseline; C2D1:cycle 2 day 1; Prog: progression.

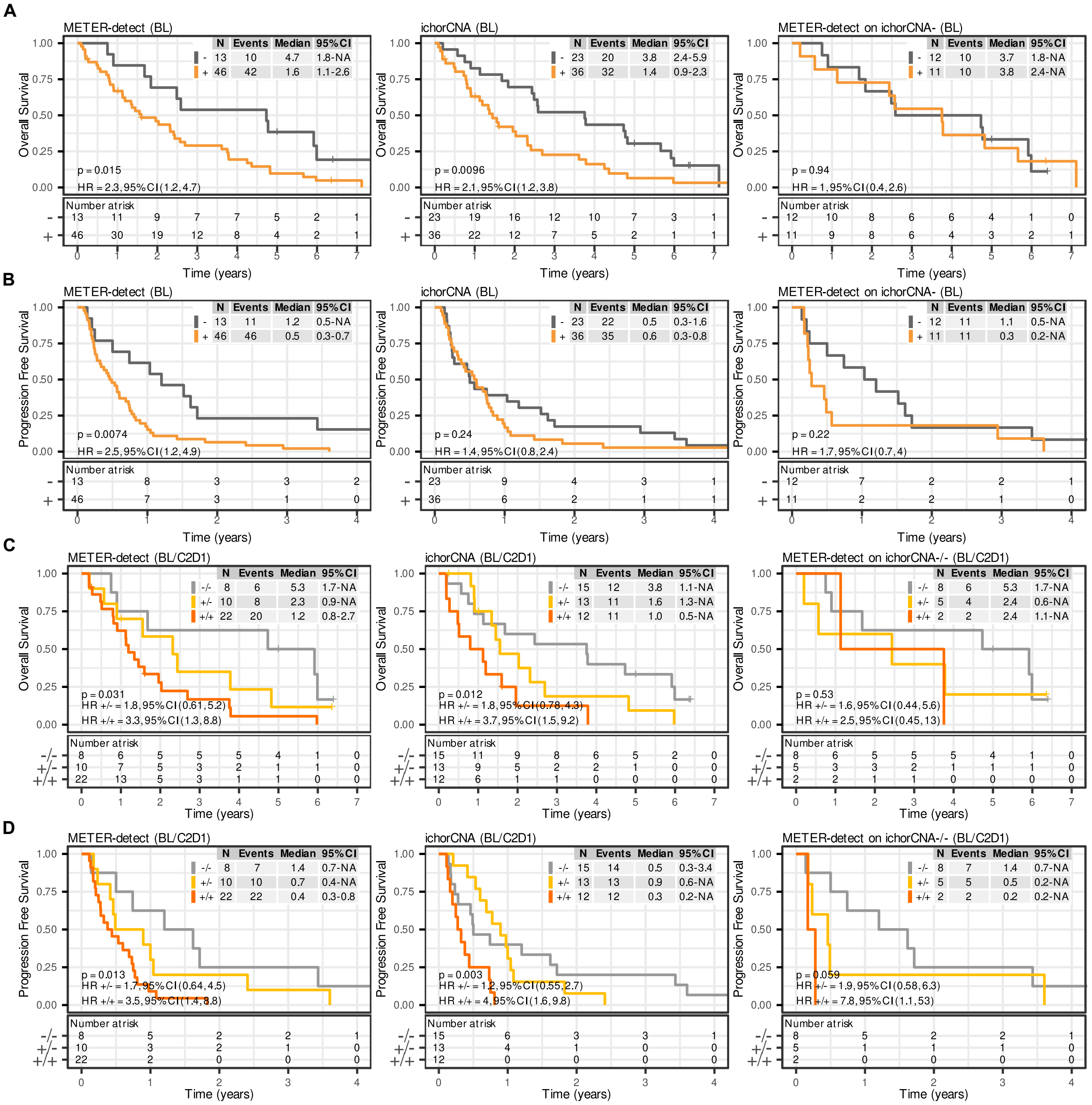


**Fig. S11: Association of patients' outcome with METER-detect classification obtained using array-based DMR detected by Rocker-meth in TCGA-BRCA dataset.** Kaplan-Meier curves showing OS (A) and PFS (B) of patients stratified according to samples' classification at BL by METER-detect with array-based iDMR (METER+ or -) and ichorCNA (ichorCNA+ or -), for all patients (N=59, left and middle panels respectively), and for ichorCNA- patients at BL (N=23, right panel) within the study cohort. (C) and (D) Kaplan-Meier curves showing (C) OS and (D) PFS of patients stratified according to samples' classification at BL and C2D1 time points (BL/C2D1) by METER-detect (METER+/+, +/- or -/-) and ichorCNA (ichorCNA +/+, +/- or -/-), for all patients having both BL and C2D1 time points (N=43, left and middle panels respectively), and for ichorCNA-/- patients at BL/C2D1 (N=15, right panel) within the study cohort. BL: pre-treatment baseline; C2D1_: cycle 2 day 1; p: Log Rank Test p-value; CI: confidence interval; HR: hazard ratio.
